# Pharmacological inhibition of LSD1 triggers myeloid differentiation by targeting GSE1, a novel oncogene in AML

**DOI:** 10.1101/2021.02.16.431315

**Authors:** Luciano Nicosia, Francesca Ludovica Boffo, Elena Ceccacci, Isabella Pallavicini, Fabio Bedin, Enrico Massignani, Roberto Ravasio, Saverio Minucci, Tiziana Bonaldi

**Author notes:** These authors contributed equally to this work (Francesca Ludovica Boffo, Elena Ceccacci).

## Abstract

The histone de-methylase LSD1 is over-expressed in haematological tumours and has emerged as a promising target for anti-cancer treatment, so that several LSD1 inhibitors are under development and testing, in pre-clinical and clinical settings. However, the complete understanding of their complex mechanism of action is still unreached. Here, we unravelled a novel mode of action of the LSD1 inhibitors MC2580 and DDP-38003, showing that they can induce differentiation of AML cells through the down-regulation of the chromatin protein GSE1. Analysis of the phenotypic effects of GSE1 depletion in NB4 cells showed a strong decrease of cell viability *in vitro* and of tumour growth *in vivo*. Mechanistically, we found that a set of genes associated with immune response and cytokine signalling pathways are up-regulated by LSD1 inhibitors through GSE1 protein reduction and that LSD1 and GSE1 co-localise at promoters of a subset of these genes at the basal state, enforcing their transcriptional silencing. Moreover, we show that LSD1 inhibitors lead to the reduced binding of GSE1 to these promoters, activating transcriptional programs that trigger myeloid differentiation. Our study offers new insights on GSE1 as a novel therapeutic target for AML.

## Introduction

By catalyzing the removal of methyl-groups from mono- and di- methylated forms of lysine 4 and lysine 9 of histone H3 (H3K4me1/me2 and H3K9me1/me2), the epigenetic enzyme Lysine-Specific histone Demethylase 1A (LSD1/KDM1A) has emerged as a major player in gene expression modulation in eukaryotes ^1,2^. In different cell types, this enzyme can act as either a transcriptional co-repressor or co-activator, depending on the distinct set of interactions established ^3^. More frequently, LSD1 is embedded in transcriptional repressive complexes, such as CoREST and NuRD ^4–6^, where different subunits modulate LSD1 activity: CoREST confers to LSD1 the ability to bind nucleosomes, directs/channels its demethylase activity towards H3K4me1/me2 and protects it from proteasomal degradation; HDACs create an hypo-acetylated chromatin environment that stimulates LSD1 catalytic activity ^6–8^. Less often, such as in the context of androgen (AR)- and oestrogen (ER)- receptor dependent transcription, LSD1 acts as transcriptional co-activator by demethylating H3K9me1/me2 and thus promoting the downstream expression of AR- and ER- target genes ^2,9^. Some proteins, such as the protein kinase C beta I (PKCbeta I) and PELP1 helps directing the LSD1 demethylase activity towards H3K9 more than H3K4 ^10,11^.

LSD1 is overexpressed in various solid and haematological tumours, where its increased levels are linked to poor prognosis ^12–15^. Various studies demonstrated LSD1 contribution to the onset and progression of Acute Myeloid Leukemia (AML), indicating that this enzyme can be a therapeutic target for treatment of different AML subtypes ^16–19^. In particular, LSD1 has been shown to stimulate the clonogenic activity of Leukemic Stem Cells (LSCs), trigger their oncogenic transcriptional programs ^16^ and also inhibit myeloid differentiation, as confirmed by the fact that LSD1 inhibition induces the activation of myeloid lineage genes, such as CD11b and CD86 ^19,20^. All these evidence have prompted the drug discovery field to develop LSD1 inhibitors as epigenetic anti-cancer drugs ^21–24^.

Because of the structural similarities of LSD1 with the monoamine oxidases (MAOs) MAO-A and MAO-B, already known inhibitors targeting MAOs have been chosen as starting scaffolds for the development of small molecules more specific/selective towards LSD1. In particular, the nonselective MAO inhibitor tranylcypromine (TCP) -which was the first compound described to efficiently inhibit LSD1 catalytic activity ^25^ - was the starting point for the design of MC2580 ^26^ and DDP-38003 ^27^, two probes that present higher potency and selectivity towards LSD1 than LSD2, MAO-A and MAO-B in in-*vitro* assays and, moreover, were shown to inhibit tumor growth and induce differentiation, when tested in murine AML blasts ^26,27^.

Various studies have helped dissecting the mechanisms of action (MoA) of these inhibitors in AML and solid tumours ^28^. Recently, a number of publications have surprisingly shown that they can trigger AML differentiation not through the expected inhibition of its catalytic activity, but by altering LSD1 binding to some of its interactors. In particular, LSD1 interaction to the transcription factors GFI1 ^19,29^ and GFI1b ^30^ was found to be strongly affected by these drugs, with the consequent effects of cell proliferation reduction and induction of myeloid differentiation in AML. These findings are particularly interesting because highlighted for the first time the role of LSD1 for the assembly of multi-protein complexes on chromatin and suggested that small molecules originally developed to target LSD1 catalytic activity can physically inhibit this scaffolding function, with therapeutic implications.

In this study, we further elaborated on our recent results on the dynamic LSD1 interactome upon its pharmacological inhibition ^29^ and focused on GSE1, whose binding to LSD1 is reduced upon cell treatment with MC2580 and DDP-38003 inhibitors, as a consequence of its diminished protein expression. Few studies have investigated the molecular and cellular function of GSE1 in cancer, so far. GSE1 has been described as an oncogene overexpressed in solid tumours, such as breast and gastric cancers, and its increased level has linked with enhanced cell proliferation, colony formation, cell migration, and invasion ^31,32^. Recently, a tumour suppressor role has been ascribed to GSE1 in neuro-epithelial stem (NES) cells ^33^. GSE1 effect in haematological malignancies and AML in particular has not been investigated yet, especially in the context of its physical and functional interaction with LSD1^34–37^.

Through the molecular and phenotypical characterization of the effect of these drugs on GSE1 expression and activity on chromatin, we provide evidence of an oncogenic role of GSE1 in AML cells and demonstrate that its drug-induced reduction enforces myeloid differentiation in AML, with relevant therapeutic implications.

## Results

### LSD1 inhibitors reduce the protein expression of GSE1 in AML

Using the Differential Enrichment analysis of Proteomics data (DEP) R software package ^38^, we re-interrogated the recently published dynamic LSD1 interactome upon treatment of NB4 Acute Promyelocytic Leukemia (APL) cells with the LSD1 inhibitor MC2580 (Fig. 1A) ^29^ and we found that, in addition to the already described GFI1, also the binding to the GSE1 protein was significantly downregulated after drug treatment (Fig 1B and 1C). We confirmed the MS results by western blot (WB) analysis, profiling GSE1 level in the LSD1 co-IP and LSD1 in the reciprocal GSE1 co-IP, in control and MC2580-treated NB4 cells. While validating the reduced interaction between LSD1 and GSE1, inspection of both experiments led to the observation that GSE1 protein was already decreased in the nuclear input of both co-IPs after LSD1 inhibition (Fig. 1D).

**Figure 1:**
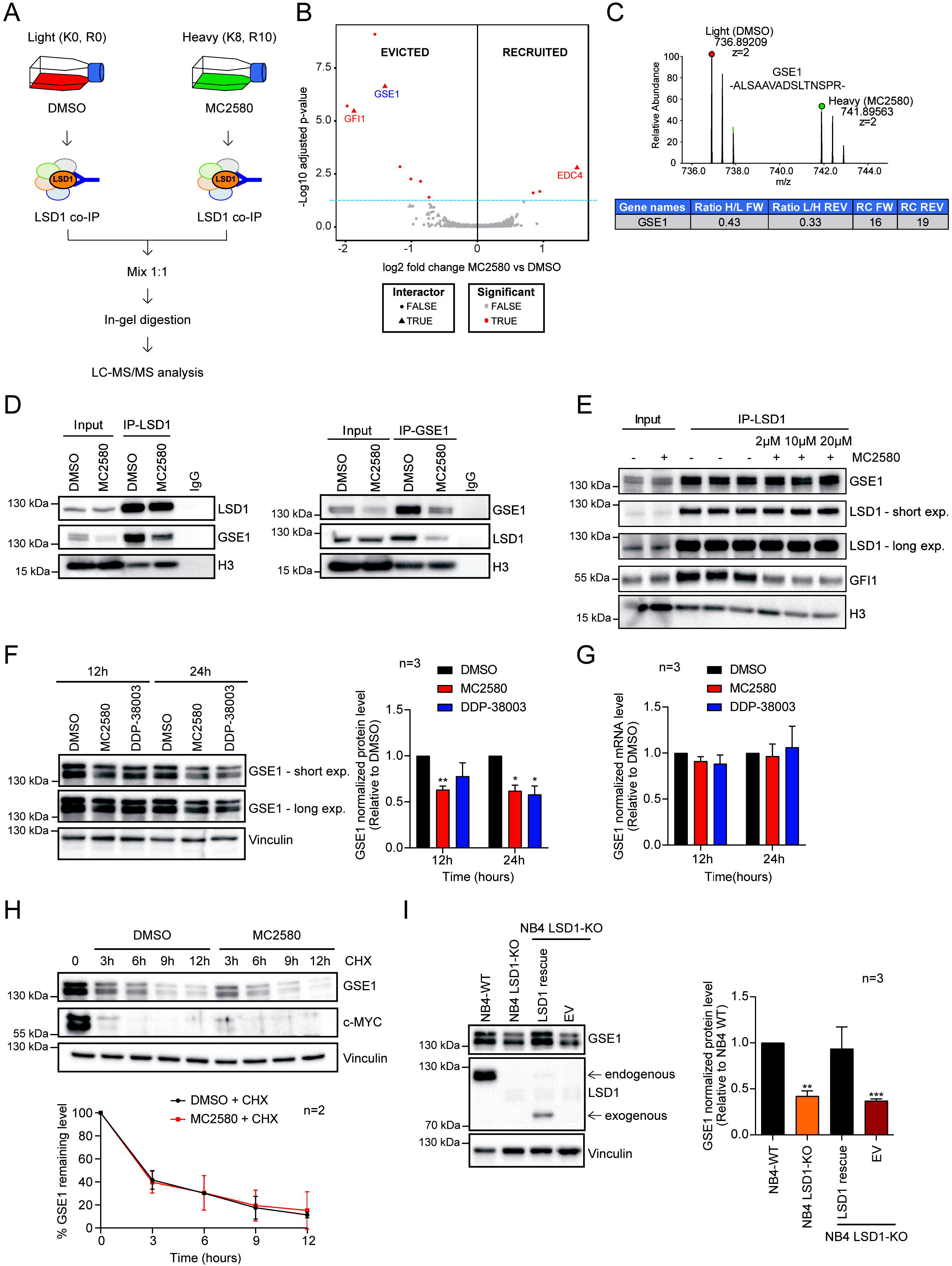
LSD1 inhibitors decrease translation of GSE1 in AML. A) Experimental design of the SILAC/LSD1 co-IP strategy setup to study the changes in LSD1 interactome upon treatment with 2 μM of MC2580, as described in ^29^. B) Volcano plot displaying the alteration in the LSD1 interactome upon MC2580 treatment. Significantly evicted and recruited LSD1 interactors are marked as red dots, while the proteins previously defined as specific LSD1 binders ^29^ are indicated by red triangles. The blue dashed line indicates the p-value threshold used to define the interactors modulated by the LSD1 inhibitors (p value < 0.05 calculated with limma ^64^). Analysis of the evicted and recruited LSD1 interactors after pharmacological treatment is performed using the Differential Enrichment analysis of Proteomics data (DEP) R software package ^38^. C) *Upper panel:* MS1 spectrum of the SILAC peak doublet for the peptide 897 -ALSAAVADSLTNSPR- 911 of GSE1 protein in the SILAC-LSD1 co-immuno-precipitated (co-IP) sample from NB4 cells treated for 24 hours with 2 μM of MC2580 (heavy peak) and DMSO-treated control cells (light peak). *Lower panel:* Table displaying the H/L (forward replicate) and L/H (reverse replicate) ratios for GSE1 in SILAC-LSD1 co-IPs of NB4 cells treated with the compound MC2580 or DMSO. The table also shows the Ratio Count (RC) of each replicate, which corresponds to the number of peptides used for SILAC-based protein quantitation. The inhibitor was added to the heavy channel in the forward replicate and the light channel in the reverse one. D) Western Blot analysis of GSE1, LSD1 and H3 in LSD1 co-IPs *(left panel)* and GSE1 co-IPs *(right panel)*, both in control and MC2580-treated cells (2 μM) for 24 hours. Rabbit IgG was used as negative control co-IP. E) Western Blot analysis of GSE1, LSD1, GFI1 and H3 in *in vitro* LSD1 co-IPs using as input NB4 nuclear cell extracts co-incubated for 6 hours with increasing doses of MC2580 (2, 10 and 20 μM) and DMSO as control. F) *Left panel:* Western Blot analysis of GSE1 in NB4 cells treated with MC2580 (2 μM), DDP-38003 (2 μM) and control DMSO for 12 and 24 hours. Vinculin was used as loading control. *Right panel:* Bar-graph displaying the quantitation results of GSE1 protein, normalized over the Vinculin, in 3 independent replicates of NB4 cells treated with MC2580 (2 μM), DDP-38003 (2 μM) and DMSO. The results are plotted as fold change (FC) of GSE1 protein level in the treated samples over control (DMSO). The chart represents mean + standard deviation (SD) (n=3 biological replicates; one sample T-test, **p value<0.01, *p-value<0.05). G) RT-qPCR analysis on GSE1 transcript in NB4 cells treated for 12 and 24 hours with either MC2580 (2 μM), DDP-38003 (2 μM) and DMSO as control. GSE1 ct values are normalized against GAPDH. The results are plotted as FC of GSE1 mRNA levels in LSD1 inhibitors-treated conditions over DMSO. Chart represents mean + SD of three (n=3) biological replicates. H) *Upper panel:* Representative Western Blot analysis of GSE1 and c-MYC in NB4 cells treated with cycloheximide (0.1 mg/ml) in combination with MC2580 (2 μM) and DMSO as control for 3, 6, 9 and 12 hours. Vinculin was used as loading control. *Lower panel:* Line-plot displaying the percentage of GSE1 protein level normalized over Vinculin in cycloheximide-treated NB4 cells, together with 2 μM of MC2580 or DMSO as control. Chart represents mean ± SD (n=2 biological replicates). I) *Left panel:* Western blot analysis of GSE1 and LSD1 in NB4 WT, NB4 LSD1-KO and LSD1-KO cells transduced with either a retroviral empty PINCO vector (EV) or a PINCO vector containing an exogenous LSD1 N-terminal truncated (172-833) form ^29^. Vinculin was used as loading control. Arrows indicate the endogenous and the exogenous LSD1. *Right panel:* Bar-graph displaying the quantitation of GSE1 protein level, normalized over Vinculin level in 3 replicates of NB4 WT, NB4 LSD1-KO and LSD1-KO cells, transduced with either an empty PINCO vector (EV) or a PINCO vector containing an exogenous LSD1 N-terminal truncated (172-833) form ^29^. In all samples the data are plotted as FC of GSE1 protein level compared to the NB4 WT, with mean + SD from n=3 biological replicates (one sample T-test, **p value<0.01, ***p-value<0.001).

Hence, differently from GFI1, we hypothesized that the reduced presence of GSE1 in the LSD1 co-IP was not due to the physical interference of their interaction by the drug, but to GSE1 diminished expression. We validated this hypothesis performing an *in vitro* interaction assay in which NB4 nuclear extract was incubated with increasing concentrations of MC2580 or DMSO as control for 6 hours, prior to carrying out the LSD1 co-IP. In line with our model, GSE1 was co-immunoprecipitated with LSD1 with the same efficiency in control- and MC2580-treated cells, while GFI1 was evicted by LSD1 after pharmacological inhibition, as previously reported ^19,29^ (Fig. 1E).

We next assessed whether the reduction of GSE1 upon LSD1 inhibition occurred only at the protein or also at the transcript level. We treated NB4 cells with MC2580 and DDP-38003 for 12 and 24 hours and measured both GSE1 mRNA and protein by real time quantitative PCR (RT-qPCR) and WB analysis, respectively: while the reduction of GSE1 protein was confirmed, peaking at 24 hours upon drug treatment (Fig. 1F), GSE1 transcript did not changed significantly (Fig. 1G), ruling out the transcriptional regulation of GSE1 upon LSD1 inhibition. This finding was corroborated by the observation that an exogenous V5-tagged form of GSE1 -cloned in two different lentiviral vectors and transduced in NB4 cells- also displayed a reduction 24 hours post-treatment with MC2580. Since the expression of the exogenous GSE1 was under the control of different promoters compared to that of the endogenous gene, this result further excludes the possibility of transcriptional inhibition of GSE1 gene by LSD1 inhibitors (Fig. S1). By assessing GSE1 levels also in other non-APL AML cell lines, such as THP-1, SKNO-1 and OCI-AML2 (Fig. S2), we confirmed that the protein down-regulation upon LSD1 inhibitors was consistent and independent from their different cytogenetic features.

In light of recent experimental evidence supporting a role of LSD1 in modulating the stability of target proteins independently from its catalytic activity ^39–42^, we next asked whether GSE1 reduction was the consequence of post-translational mechanisms associated with protein destabilization. First, we assessed GSE1 protein level at different time points upon treatment with cycloheximide (CHX), a translational elongation inhibitor used to monitor protein stability, and observed a strong reduction 12 hours post-treatment, while c-MYC was efficiently degraded 3 hours post CHX treatment, as already described ^43^ (Fig. S3). Then, we profiled GSE1 protein level at 3,6, 9 and 12 hours upon CHX treatment in the presence or absence of MC2580 and observed that GSE1 protein stability was not altered when LSD1 was pharmacologically inhibited, suggesting that the diminished level of GSE1 is not due to the decreased stability or enhanced degradation of the protein, but likely to translation impairment (Fig. 1H, Fig. S4).

Pharmacological data were corroborated by the analysis of LSD1 knock-out (KO) cells, which showed diminished GSE1 protein compared to NB4 wild-type (WT) cells (Fig. 1I). Last, when we transduced LSD1 KO cells with a vector re-expressing an exogenous WT form of LSD1 ^29^, we observed GSE1 protein re-established to a level significantly higher than in the cells transduced with an empty vector (EV) (Fig. 1I). This result specifically links GSE1 protein reduction to the inhibition or depletion of LSD1, excluding off-targets effects induced by the drugs.

### GSE1 is an oncogene important for viability and *in vivo* tumour growth of NB4 cells

We then set to explore the phenotypic effects of GSE1 down-regulation in AML. To do it, we silenced GSE1 in NB4 cells by RNA interference, using two short hairpin RNAs (shRNAs) that displayed different efficiency in depleting GSE1, with shB2 being stronger than shA1 (Fig. 2A). We observed a robust reduction of cell viability that appeared dependent on GSE1 level. Alongside with reduced cell growth, cell death measured by trypan blue staining also increased with time and correlated positively with silencing efficiency (Fig. 2B and 2C). This result was corroborated by the detection of the cleaved caspase-3, a marker of apoptosis, in both shA1- and shB2- transduced cells 72 hours post-infection (Fig. 2D). The link between GSE1 KD and apoptosis was further confirmed in the shB2-transduced cells by measuring the reduction of total (not cleaved) caspase-3, which indicated that, in these cells, the majority of the enzyme was in its active form already 72 hours post-infection (Fig. 2D). Together, these experiments demonstrated that GSE1 depletion impairs NB4 cell growth and induces apoptosis.

**Figure 2:**
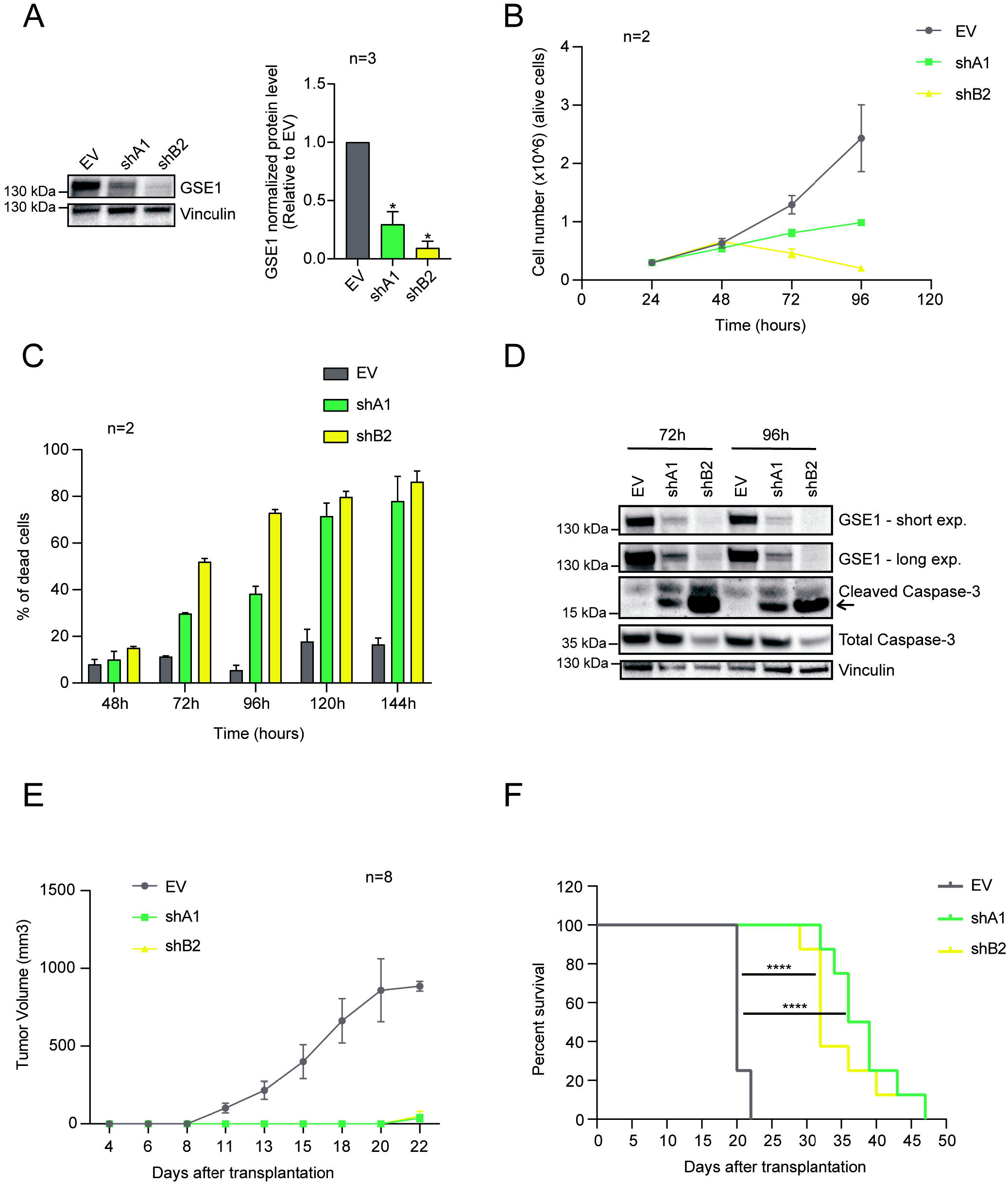
GSE1 knock-down (KD) affects NB4 cell viability and *in vivo* tumour growth in NOD/SCID gamma (NSG) mice. A) *Left panel:* Western blot analysis of GSE1 protein in NB4 cells transduced with either an empty lentiviral pLKO.1 puro vector (EV) or pLKO.1 puro containing the shRNA constructs targeting GSE1 (shA1 and shB2) for 72 hours. Vinculin was used as loading control. *Right panel:* Bar-graph displaying the quantitation of GSE1 protein level normalized over the Vinculin in three independent replicates (n=3), whereby the results are plotted as fold change (FC) of GSE1 level in the knock-down (KD) samples over the control EV. Chart represents mean + standard deviation (SD) (n=3 biological replicates; one sample T-test, *p value<0.05). B) Growth curve of NB4 cells transduced with either an empty pLKO.1 puro vector (EV) or pLKO.1 puro containing the shA1 and shB2 targeting GSE1. Graph represents mean ± SD from two biological replicates (n=2). C) Bar-graph displaying the percentage of dead cells in control EV and GSE1 KD cells. The analysis is performed after 48, 72, 96, 120 and 144 hours of infection. Cell viability was measured by trypan blue staining. Chart represents mean + SD from two (n=2) biological replicates. D) Western Blot analysis of GSE1, cleaved caspase-3 and total caspase-3 in cells transduced with either the control pLKO.1 puro (EV) or pLKO.1 containing the two shRNAs targeting GSE1 (shA1 and shB2) for 72 and 96 hours. Vinculin was used as loading control. The arrow indicates the cleaved form of the caspase-3. E) Tumour growth curve in NSG mice transplanted subcutaneously with NB4 cells transduced 24 hours earlier with either an empty pLKO.1 puro or pLKO.1 puro with the insert of the shRNAs (shA1 and shB2) targeting GSE1. Graph represents mean ± SD of the tumour volume in each condition (n=8, 8 distinct mice for each group). F) Kaplan-Meier survival curve of mice transplanted with NB4 cells previously transduced with either an empty pLKO.1 vector or pLKO.1 containing shA1 or shB2 targeting GSE1 (n=8 for each group; Log-rank Mantel-Cox test, ****p-value<0.0001).

We also assessed the effect of GSE1 downregulation on cell cycle progression by measuring the DNA content of cells with Propidium Iodide (PI), at early time points upon infection: we observed a 10%-20% increase of the proportion of cells in G1-phase, mirrored by a corresponding decrease in the percentage of cells in S-phase, in GSE1 KD cells compared to control EV-transduced ones (Fig. S5).

Based on these results *in vitro*, we tested the effect of GSE1 depletion on tumor growth *in vivo:* 24 hours post transduction with shA1, shB2 and control EV (Fig. S6), NB4 cells were injected subcutaneously in NSG mice and tumor growth was measured at time intervals, until control EV-injected mice were sacrificed when the tumor reached a maximal diameter of about 15 mm. GSE1 KD strongly affected tumor growth, as indicated by the fact that mice injected with GSE1 KD cells presented palpable tumors only at day 22 after transplantation (Fig. 2E), when almost all control mice had already been sacrificed. Furthermore, the survival curve showed that mice transplanted with EV-transduced NB4 cells died between day 20 and day 22, whereas mice transplanted with GSE1-depleted cells had a prolonged lifespan, with median survival time of 38 days (Fig. 2F). These *in vivo* results confirm the *in vitro* data and indicate that GSE1 is a relevant oncogene in AML.

### Reducing GSE1 protein level in AML, LSD1 inhibitors promote the activation of cytokine-mediated signaling and immune response pathways

Next, we set to investigate the molecular mechanisms underpinning the phenotypic effects observed. Since GSE1 is a subunit of different transcriptional regulatory complexes, we decided to assess the impact of GSE1 depletion on the NB4 transcriptome, carrying out RNA-sequencing (RNA-seq) analysis of cells infected with either the two GSE1- shRNAs, or the EV as control. The analysis was carried out at 48 hours post-infection, a time point when cell death is still negligible (Fig. 2C). Upon GSE1 KD, 720 genes were up-regulated and 131 down-regulated in shA1- infected cells, and 999 were up-regulated and 521 down-regulated genes in shB2- transduced NB4 (Fig. 3A, Table S1). This is in line with the evidence that GSE1 is mainly associated with co-repressor complexes, such as the HDAC2 and BHC complexes ^35,37^ Intersecting the differentially expressed genes (DEG) in common between the two KD conditions, we obtained 422 common up-regulated and 47 common down-regulated genes (Fig. 3B). Gene ontology (GO) analysis of the up-regulated gene group revealed an enrichment of terms related to immune and inflammatory responses and cell proliferation, while no significant enrichment of specific biological process occurred in the down-regulated gene group (Table S2). In particular, using the Revigo Web server ^44^, we found the enrichment of the biological process (BP) terms “cytokine mediated signalling”, “negative regulation of viral genome replication”, “regulation of cell migration”, “extracellular matrix organization” and “regulation of cell proliferation”. Within the term “negative regulation of viral genome replication” we found biological processes associated with both immune response - like “neutrophil mediated immunity” and “negative regulation of leukocyte mediated cytotoxicity”- and myeloid differentiation, such as “regulation of monocyte differentiation”. Instead, the term “cytokine mediated signalling” included several BP sub-terms linked to inflammation, like “regulation of I-kappaB kinase/NF-kappaB signalling” (Fig. 3C). Similar results were obtained when we analysed the upregulated gene sets with Reactome, to highlight the significant enriched biological pathways ^45^ (Fig. S7).

**Figure 3:**
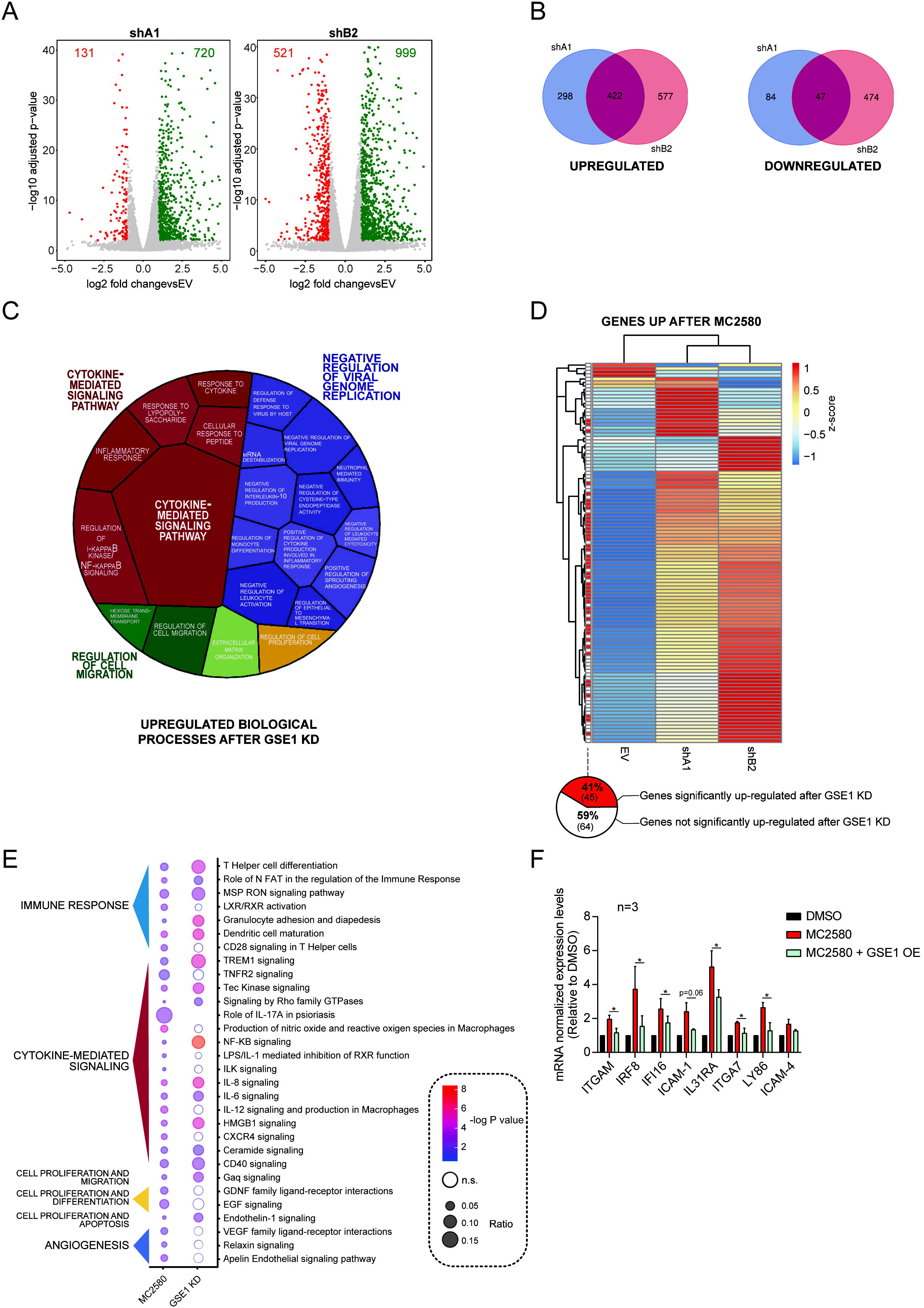
GSE1 down-regulation elicited by LSD1 inhibitors induces transcription of genes associated with cytokine-mediated signalling and immune response pathways. A) Volcano plot displaying up- and down- regulated genes upon 48 hours transduction with pLKO.1 vector containing shA1 and shB2 inserts. The x-axis shows the log2 fold change (FC) values of each gene in the shRNAs-transduced cells compared to control empty vector (EV)-transduced, while y-axis displays the -log_10_ adjusted p-values (p-adj). FC value is calculated with DEseq2 program ^53^, using two biological replicates for each condition (n=2). B) Venn diagrams with number of individual and overlapping up-regulated and down-regulated genes identified 48 hours after transduction with shA1 and shB2 constructs. C) Voronoi tree-map displaying the statistically significant GO biological process (BP) terms associated with the common 422 up-regulated genes upon GSE1 KD with the two shRNA constructs. GO analysis and calculation of the statistically significant BP was performed through EnrichR ^54^ (adjusted p-value < 0.05), and then BP were grouped using Revigo ^44^. Voronoi plot includes the BP terms mapped to a high hierarchical level. The tassel size corresponds to the -log_10_ p-value of the enrichment while the colour intensity to the number of genes belonging to each category. D) Unsupervised hierarchical clustering of all significantly up-regulated genes upon MC2580 treatment (p-adj < 0.01, log2 FC > 1). Heatmap displays the RNA-seq expression z-scores of these genes 48 hours after transduction with pLKO.1 puro containing shA1, shB2 or the EV as control in NB4 cells. Each row in the heatmap represents an individual gene. Z-score values of each gene are plotted as the average of two independent biological replicates, for each condition (n=2). Genes significantly up-regulated by both shA1 and shB2 (p-adj < 0.01, log2 FC > 1) are flagged in red. Below is shown the pie-chart indicating the percentage of the 109 genes up-regulated by MC2580 that are also significantly induced by GSE1 KD with both shRNAs. E) Bubble-plot displaying the statistically significant IPA pathways (-log10 p-value > 2) obtained from the analysis of the differentially expressed genes (DEG) after MC2580 treatment and comparison with the IPA results achieved from the DEG upon GSE1 KD. The bubble size reflects the number of genes belonging to that pathway, while colours indicate the -log_10_ p-value associated with each IPA term. Non-significant IPA pathway terms are coloured in white. IPA pathways are clustered manually, according to specific biological processes. Some IPA pathway terms are not displayed in the current figure panel, but the complete list is in the Table S4. F) RT-qPCR analysis on a panel of genes up-regulated by both LSD1 inhibitors and GSE1 KD, in control and GSE1 over-expressing (OE) NB4 cells treated with 2 μM of MC2580 treatment. GSE1 OE cells are generated by lentiviral transduction with the pLEX_307 vector containing the GSE1 coding sequence (NM_001134473.3), while control cells are transduced with the empty pLEX_307 (pLEX_307 EV). Ct values for each gene are normalized against GAPDH. The results are plotted as FC of the mRNA levels in the treated conditions relative to the control DMSO. Bar-graph represents mean + standard deviation (SD) from three (n=3) biological replicates; paired T-test, *p-value<0.05).

The transcriptomic data on the one hand confirmed the phenotypic results obtained by the cellular assays carried out upon GSE1 depletion, such as the activation of genes involved in the regulation of cell proliferation, on the other hand highlighted the possible role of this protein in other processes, like myeloid differentiation and inflammatory response.

In light of the observed link between LSD1 pharmacological inhibition and GSE1 down-regulation, we then compared the transcriptomic changes observed upon GSE1 depletion with the RNA-seq data of NB4 cells treated with MC2580, in order to unravel possible overlapping transcriptional programs that may suggest the presence of molecular pathways activated by LSD1 inhibitors, in dependence of GSE1. Compared to GSE1 KD, MC2580 induced significantly lower transcriptional changes, with only 109 significantly up-regulated and 3 down-regulated genes, respectively (Fig. S8, Table S3). This is in line with the stronger cellular effects observed upon GSE1 down-regulation compared to those elicited by LSD1 pharmacological inhibition. Under the hypothesis that a subset of genes may be transcriptionally induced by the drug through GSE1 downregulation, we first assessed the expression levels of the 109 genes up-regulated by MC2580 within the GSE1 KD cell transcriptome. Most of the genes modulated by MC2580 also showed an increasing trend upon GSE1 depletion; in particular, 40% of them were significantly up-regulated after infection with both shRNAs (Fig. 3D). Given this overlap, we assessed through Ingenuity Pathway Analysis (IPA) whether the molecular pathways activated by LSD1 inhibition were the same as those stimulated by GSE1 depletion: 55% of the pathways triggered by the drug were also significantly activated by GSE1 depletion and the majority were associated with “cytokine-mediated signaling” and “immune response” GO terms (Fig. 3E, Table S4). We confirmed the up-regulation of some of the genes belonging to these biological processes by RT-qPCR upon both MC2580/DDP-38003 treatment (Fig. S9A) and GSE1 KD (Fig. S9B).

These results suggest that LSD1 inhibitors upregulate genes belonging to cytokine-mediated signalling and immune response pathways via GSE1 down-regulation. This hypothesis was confirmed by profiling the expression of a panel of genes involved in these processes in NB4 cells upon MC2580 treatment, in combination or not with GSE1 over-expression. Observing that the up-regulation of these genes upon LSD1 inhibition was significantly reduced when GSE1 was over-expressed (Fig. 3F), we propose a novel function of GSE1 as an important co-repressor of genes linked to inflammatory- and immune- response pathways and suggest the use of these inhibitors to unlock them through GSE1 targeting.

### LSD1 pharmacological inhibition reduces GSE1 association to LSD1-bound promoters of genes involved in immune response and inflammatory pathways

Transcriptomic analysis suggested that the pharmacological inhibition of LSD1 leads to the GSE1-dependent activation of specific transcriptional programs. Since GSE1 is a chromatin-associated factor, we decided to profile the effect of its reduction on chromatin by performing Chromatin Immuno-Precipitation (ChIP). ChIP experiments were carried out in NB4 cells upon expression of the V5-tagged form of GSE1, whereby the V5-tag was used as bait for affinity enrichment of bound chromatin, to overcome the problem of the unavailability of ChIP-grade antibodies against GSE1. As the drug induces down-regulation of the exogenous V5-GSE1 to a similar extent to the endogenous protein (Fig. S1), we reasoned that ChIP-seq analysis of the V5-tagged isoform could be a good proxy to assess the effects of LSD1 inhibitors on GSE1 genomic localization.

ChIP-seq analysis proved that GSE1 binding to chromatin was globally reduced upon drug treatment, with the number and overall intensity of assigned peaks almost halved in treated cells (Fig. 4A and 4B). By inspection of the genomic distribution of the assigned peaks in both functional states, we found that at basal state GSE1 mainly localizes at distal intergenic (~ 25%), intronic (~ 35%) and promoter (~ 25%) regions, while upon LSD1 inhibition its binding slightly increases at distal intergenic and intronic regions and decreases at the promoters (Fig. 4C).

**Figure 4:**
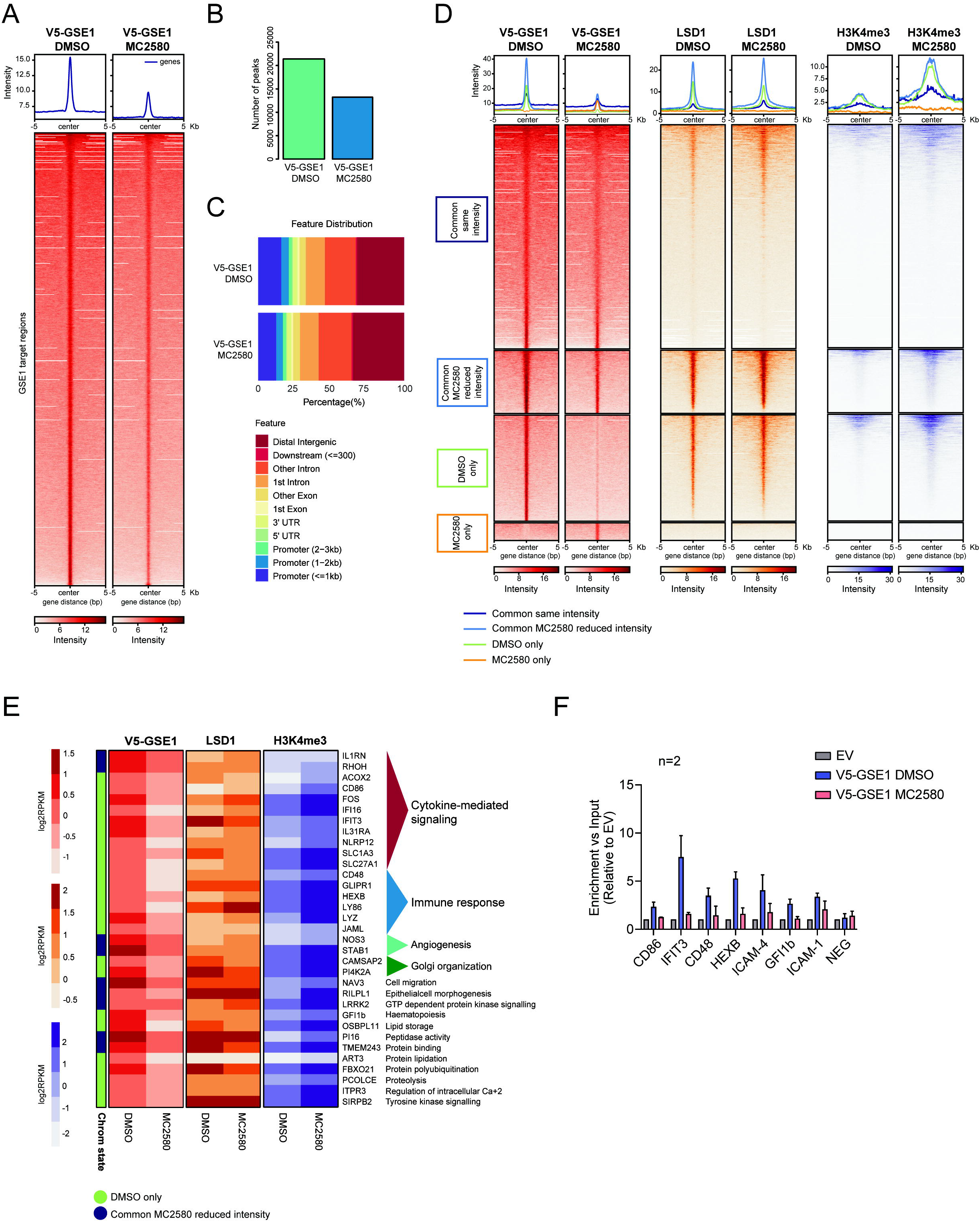
MC2580 decreases GSE1 localization on the LSD1-bound promoters of genes involved in cytokine-mediated signalling and immune response pathways. A) Heatmap representing the normalized V5-GSE1 ChIP-seq intensities ±5 kb around the center of the GSE1 target loci, at basal state and after 24 hours treatment with MC2580 (2 μM). V5-GSE1 ChIP-seq was performed in NB4 GSE1 OE cells produced by lentiviral transduction of the pLEX_307 construct containing the GSE1 coding sequence (NM_001134473.3) (pLEX_307 GSE1), while the negative control ChIP was carried out in cells transduced with the empty pLEX_307 (pLEX_307 EV). B) Bar-graph displaying the number of peaks extrapolated from the V5-GSE1 ChIP-seq in DMSO and MC2580-treated cells. C) Genome annotation of the V5-GSE1 ChIP-seq peaks in control DMSO and LSD1-inhibited NB4 cells. D) Heatmap displaying the normalized V5-GSE1, LSD1 and H3K4me3 ChIP-seq intensities ±5 kb around the center of the GSE1 target loci after 24 hours treatment with MC2580 or DMSO as control. The heatmap is grouped in different subgroups, according to the presence or absence or differential intensity of the GSE1 binding regions in the two conditions. The subset of “Common GSE1 targets with reduced intensity in DMSO” contains only 32 regions and is not displayed in the figure. The LSD1 and H3K4me3 ChIP-seq used to generate the heatmap were published in ^29^. E) Heatmap displaying the V5-GSE1, LSD1 and H3K4me3 ChIP-seq intensities at the promoters of genes up-regulated after LSD1 inhibition, which show reduced binding of V5-GSE1 in MC2580 treated-cells. Each row in the heatmap represents an individual gene. For the V5-GSE1 ChIP-seq, reads intensities in regions identified as peaks and annotated to each gene are plotted as log2 RPKM. Instead for the LSD1 and H3K4me3 ChIP-seq, reads coverage (log2 RPKM) at promoter (+/- 2.5 kb distance from the TSS) were represented. For each gene its chromatin state is shown as referred in the Fig. 4D. F) ChIP-qPCR profiling of a panel of GSE1-bound promoters of genes significantly up-regulated by LSD1 inhibition in DMSO and MC2580 (2 μM)-treated NB4 GSE1 OE cells using the pLEX_307 GSE1 construct. V5 ChIP in NB4 cells transduced with the pLEX_307 EV was used as negative control ChIP. NEG corresponds to a negative control region. Data are normalized over the respective Input and displayed as fold change (FC) over the control EV. Bar-graph represents mean + standard deviation (SD) from two (n=2) biological replicates.

We then grouped the genomic regions bound by GSE1 in four different categories: “Regions bound by GSE1 similarly in both DMSO- and MC2580-treated cells”, “GSE1- bound regions that display reduced binding upon MC2580”, “regions bound by GSE1 only in DMSO condition” and “GSE1- bound regions only upon MC2580” (Table S5). We profiled by ChIP-seq the localisation of LSD1 and different histone post-translational modifications (PTMs) at these four genomic regions and found that LSD1 and all histone PTMs analysed co-localized with V5-GSE1 in all the genomic regions, except from those bound exclusively by GSE1 upon drug treatment. Furthermore, LSD1 binding at GSE1 regions was not affected by the inhibitor, while histone PTMs associated with active transcription (mainly H3K4me3 and slightly H3K4me2) were enriched, particularly at those loci where GSE1 binding was reduced/abolished by the compound (“GSE1- bound regions that display reduced binding upon MC2580” and “regions bound by GSE1 only in DMSO condition”) (Fig. 4D, Fig. S10). This suggests that the decreased association of GSE1 to chromatin may be linked to a chromatin state more prone to transcriptional activation. Moreover, in light of the reduced localization of GSE1 at promoters upon MC2580 and of the increased H3K4me3 level observed at the same regions (Fig. 4D, Fig. S11), we hypothesized that the decreased GSE1 association could directly mediate the transcriptional changes measured upon LSD1 inhibition. To confirm this hypothesis, we assessed the presence of promoters of the genes up-regulated by MC2580 within the subsets of genomic regions where GSE1 binding was diminished by the drug and found several promoters of genes associated with cytokine-mediated signalling and immune response, like IFI16, IL31RA, CD86, FOS, CD48. LSD1 co-localized with GSE1 at these promoters and its binding was not altered by the drug, while the H3K4me3 level increased (Fig. 4E). We validated the ChIP-seq results by ChIP-qPCR for some of these promoters. With ChIP-qPCR we also detected a reduced binding of GSE1 at the promoters of ICAM-1 and ICAM-4 upon MC2580, other two interesting genes involved in cytokine signalling pathways that were up-regulated by LSD1 inhibition (Fig. 4F). The ChIP data corroborate the transcriptomic results and provide a mechanistic insight about the effect of LSD1 inhibitors on GSE1 localisation and activity at chromatin regulatory regions.

### GSE1 down-regulation induced by LSD1 inhibition triggers myeloid differentiation

Pharmacological inhibition of LSD1 represents a promising epigenetic approach for AML treatment through the release of the differentiation block and the induction of differentiation processes in leukemic blast cells ^17,19^. In line with this, our transcriptomic data demonstrated that the majority of the DEGs induced both by MC2580 and GSE1 depletion were involved in haematological system development and haematopoiesis (Fig. S12). Thus, we hypothesized that the LSD1-dependent reduction of GSE1 on chromatin might be phenotypically linked to the induction of myeloid differentiation. To test this hypothesis, first we profiled by flow cytometry the expression of the cell surface differentiation marker CD11b in GSE1 KD NB4 cells, observing 4- and 8- fold increase in the percentage of cells expressing this marker 48 hours post transduction with shB2 and shA1 constructs, respectively (Fig. 5A). Induction of CD11b upon GSE1 KD was detected also in THP-1 monocytic cells, where we could also assess CD14, the monocyte differentiation antigen typically up-regulated upon a differentiation stimulus: 4- to 6- fold increase of CD14 was observed in GSE1 KD cells compared to the control. Furthermore, the majority of CD14- positive cells were also positive to CD11b (Fig. 5B).

**Figure 5:**
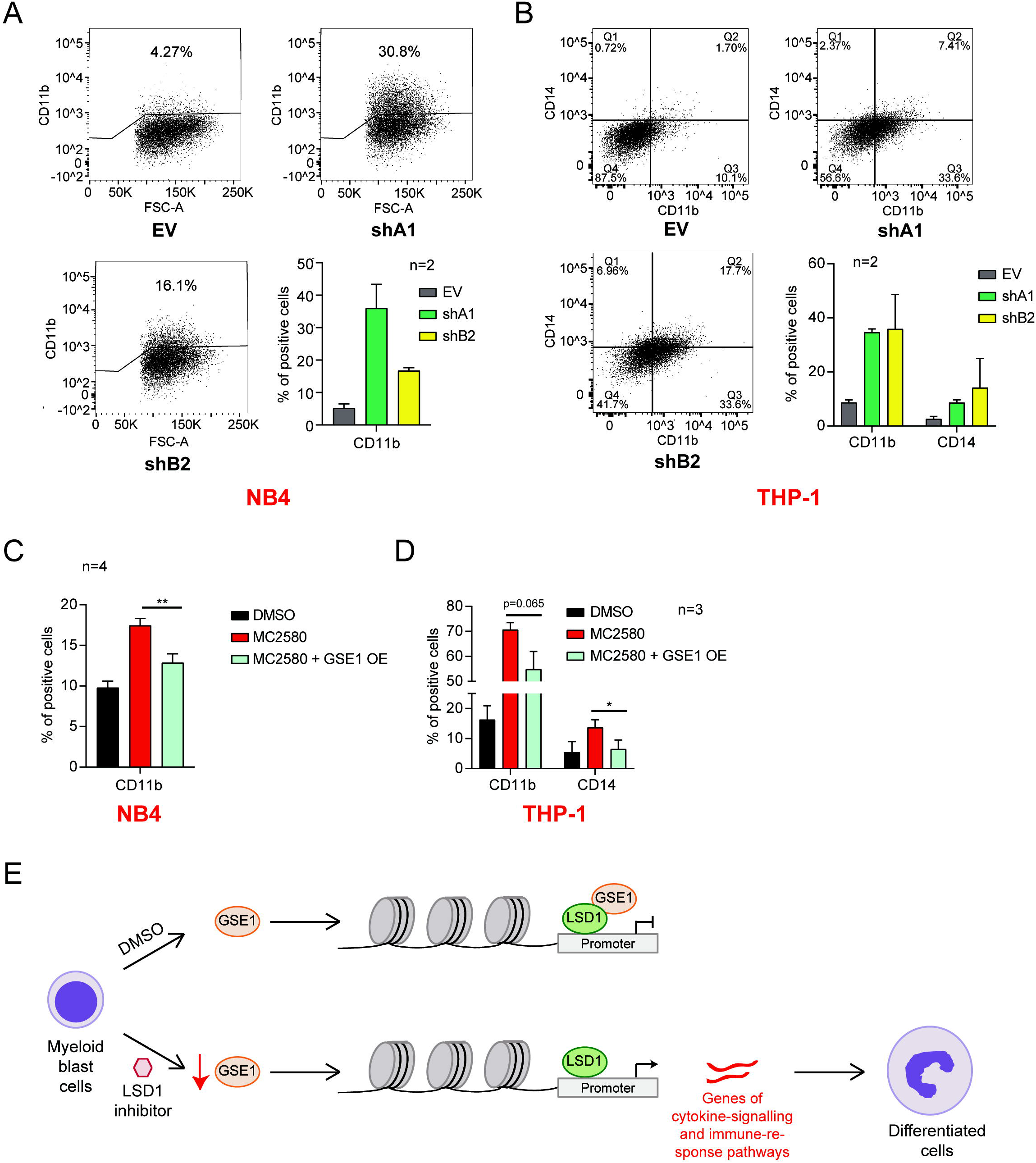
GSE1 down-regulation contributes to the myeloid differentiation re-activation enforced by LSD1 inhibition. A) Percentage of NB4 CD11b-positive cells measured by flow cytometry 48 hours post transduction with the shA1 and shB2 cloned in pLKO.1 puro, or with the empty pLKO.1 puro vector (EV) used as negative control. Dot plots show the results obtained in one of the two biological replicates of the experiment. Bar-chart represents mean + standard deviation (SD) (n=2 biological replicates). B) Percentage of CD11b- and CD14- positive THP-1 cells, assessed by flow cytometry analysis 48 hours upon transduction with pLKO.1 puro vector containing shA1, shB2 or EV as negative control. Dot plots display the data of one of the two biological replicates of the experiment. Bar-graph shows mean + SD (n=2 biological replicates). C) Percentage of CD11b-positive cells evaluated by flow cytometry in control (EV) and GSE1 OE cells, treated for 24 hours with MC2580 (2 μM). NB4 GSE1 OE are produced by transducing the pLEX_307 vector with the GSE1 coding sequence (NM_001134473.3), while control cells are transduced with the pLEX_307 EV. Chart represents mean + SD (n=4 biological replicates; paired T-test, **p-value<0.01). D) Flow cytometry analysis of the percentage of CD11b- and CD14-positive cells in EV and GSE1 OE THP-1 cells treated for 24 hours with MC2580. THP-1 GSE1 OE are generated by lentiviral transduction of the pLEX_307 vector including GSE1 coding sequence (NM_001134473.3), while control cells are transduced with the pLEX_307 EV. Chart represents mean + SD (n=3 biological replicates; paired T-test, **p-value<0.01). E) Model of the molecular effects associated with the down-regulation of GSE1 induced by LSD1 pharmacological inhibition.

Then, we asked whether GSE1 reduction induced by LSD1 inhibition was important to trigger myeloid differentiation by measuring with flow cytometry CD11b expression at 24 hours post MC2580 treatment, in wild type (EV-infected) NB4 cells and in the presence of GSE1 over-expression (OE). The observation that the increased CD11b level upon LSD1 inhibition was significantly reduced when GSE1 was overexpressed suggests that the differentiation process elicited by MC2580 also depends on GSE1 protein level (Fig. 5C). We also analysed the expression of CD11b and CD14 in control (EV) THP-1 cells and in GSE1 overexpressing ones and confirmed the same results observed in NB4 (Fig. 5D). Interestingly, the overexpression of GSE1 in NB4 also led to the increase of the subpopulation of cells expressing the stemness marker cKIT. This indicates that GSE1 not only prevents myeloid differentiation but also favours the maintenance of a stem phenotype, at least in this cell line (Fig. S13 and S14).

Altogether the data collected led to the elaboration of a model whereby the inhibition of LSD1 with MC2580 and DDP-38003 reduces GSE1 protein level, leading to its reduced binding to chromatin and in particular to LSD1-target promoter regions regulating the expression of genes linked with cytokine-signalling and immune response pathways. The consequent up-regulation of these transcriptional programs enforces myeloid differentiation in leukemic cells (Fig. 5E).

## Discussion

In this study, we show for the first time that the LSD1 inhibitors MC2580 and DDP-38003 can elicit myeloid differentiation in AML through the down-regulation of GSE1 protein, a poorly explored LSD1 interactor. Our data add another layer of information on the mechanism of action of these drugs in leukaemia, beyond the already known inhibitory effects on the lysine histone demethylase activity ^20^ and on LSD1-GFI1 interaction ^19,29^. Furthermore, our results unveil the oncogenic role of GSE1 in AML.

Our data suggest that the drugs in use most likely affect GSE1 translation rather than transcription or stability, even if the details of the mechanism remain to be elucidated. One possibility is that the decrease of GSE1 protein level may be caused by the drug-induced upregulation of some miRNA targeting this gene. This hypothesis is supported by the evidence that various miRNAs can modulate GSE1 expression ^31,32,46^. Alternatively, GSE1 downregulation could be caused by the block of its translational initiation ^47^, in line with recent data showing that -in some cell lines- LSD1 inhibitors activate the mTOR signalling cascade ^48,49^, which directly affects protein synthesis ^50^. Future investigations will allow dissecting mechanistically how LSD1 inhibitors modulate GSE1 translation.

The RNA-seq data show that in NB4 cells GSE1 depletion induces more extensive transcriptional changes than those caused by LSD1 inhibitors. In line with this, also the phenotypic effects elicited upon GSE1 KD are more pronounced than those observed upon treatment with LSD1 inhibitors, which consist in a mild reduction of cell viability and colony-forming ability in liquid and semi-solid culture, respectively ^29^. These molecular and phenotypic differences can be explained in light of the significantly different reduction of GSE1 protein level caused by these two perturbations, with MC2580/DDP-38003 leading to a milder GSE1 down-regulation (around 40%) than the shA1 and shB2 constructs (around 80%-90%). The detected dose-dependency of the phenotypic effects to GSE1 reduction induced by the two shRNA constructs used in this study corroborates this hypothesis.

The evidence that cellular levels of GSE1 are critical for NB4 cell viability is particularly intriguing as it suggests that this oncogene can represent a new target for AML treatment.

From a molecular standpoint, the transcriptomic data collected in this study indicate that the induction of genes involved in cytokine-signalling and immune response pathways by LSD1 inhibitors is GSE1-dependent. Specifically, we propose that this transcriptional response is mediated by the reduction of GSE1 on chromatin and in particular, among the different genomic regions bound by GSE1, we focused on promoters -most of them bound by also LSD1- because of the possibility to directly link the effects of GSE1 binding variation to the expression of the associated genes, by the intersection of ChIP-seq and RNA-seq data. It shall be noted, however, that the ChIP-seq data also showed the binding of GSE1 to distal intergenic and intronic regions and the overall co-localization with H3K4me1 and H3K27ac, two histone modifications traditionally associated with enhancers. This observation points towards the presence of GSE1 at other *cis-*regulatory elements, so that its reduced association to chromatin after drug treatment could cause more global transcriptional effects, still to be described.

In summary, by describing for the first time the molecular and cellular implications of GSE1 modulation in AML, this study paves the way to the thorough assessment of the role of this chromatin factor in haematological malignancies and of the possibility of targeting it with LSD1 inhibitors to trigger myeloid differentiation for AML treatment.

## Materials and Methods

### Cell culture

NB4 and THP-1 cell lines were grown in RPMI plus 10% of foetal bovine serum (FBS), 2 mM glutamine (Glu) and 1% Penicillin/Streptomycin (P/S). SKNO-1 cells were cultured in the same conditions, with the addition of 10 ng/ml GM-CSF. OCI-AML2 cell line was grown in 80% alpha-MEM supplemented with 20% FBS, 2 mM Glu and 1% P/S. Cultures were maintained in a humidified tissue culture incubator at 37°C in 5% CO2.

### Compounds

The LSD1 inhibitors MC2580 and DDP-38003 were provided by the Department of Drug Chemistry and Technologies of the Sapienza University of Rome (Italy) and the Experimental Therapeutic Unit of the IFOM-IEO Campus, respectively ^26,27^. Cycloheximide was purchased from Sigma Aldrich (C7698).

### RNA-sequencing (RNA-seq) and data analysis

mRNA-sequencing (mRNA-seq) libraries were prepared with the TruSeq RNA Sample Preparation v2 kit (RS-122-2002, Illumina) according to the manufacturer’s protocol, starting from 500 ng of total RNA per sample. Sequencing was performed using the NovaSeq 6000 (Illumina) instrument. Raw reads were mapped to the human reference genome hg38 using STAR aligner ^51^ and quantified through the *rsem-calculate-expression* function of the RSEM package ^52^. Differentially expressed genes (DEG) were determined with the DEseq2 package ^53^, as follows: genes with an adjusted p-value lower than 0.01 and a log2 Fold Change (FC) greater than 1 and smaller than −1 (log2 FC < −1 and > 1) were considered as upregulated and downregulated, respectively. GO analysis of the enriched biological processes (BP) was carried out with the EnrichR software ^54^, while biological pathway analysis was executed using the Reactome database ^45^ contained within EnrichR software and the Ingenuity Pathway Analysis (IPA) (QIAGEN Inc., https://www.qiagenbioinformatics.com/products/ingenuitypathway-analysis). Significant BP and Reactome terms had an adjusted p-value < 0.05, while significant IPA pathways had a –log(p-value) > 2. Voronoi plot of the GO was generated using the R-package voronoiTreemap (https://github.com/uRosConf/voronoiTreemap).

### Lentiviral and retroviral constructs for exogenous protein expression

NB4 LSD1 KO cells and NB4 LSD1 KO cells transduced with the LSD1 N-terminal truncated (172-833) form or the empty PINCO vector were generated as previously described in ^29^. To knock-down (KD) GSE1, short hairpin RNA (shRNA) constructs targeting the protein were cloned into the pLKO.1 puro expression vector. The primers used for shRNA production and cloning were the following:

1. shA1:

a. FW 5’-CCGGGAACTCACCTTGACGTCAATGCTCGAGCATTGACGTCAAGGTGAGTTCTTTTTG-3’
b. REV 5’-AATTCAAAAAGAACTCACCTTGACGTCAATGCTCGAGCATTGACGTCAAGGTGAGTTC-3’
2. shB2:

a. FW 5’-CCGGCTGAGCATGCTTCACTATATCCTCGAGGATATAGTGAAGCATGCTCAGTTTTTG-3’
b. REV 5’-AATTCAAAAACTGAGCATGCTTCACTATATCCTCGAGGATATAGTGAAGCATGCTCAG-3’

To over-express GSE1, its coding sequence transcript (GenBankTM accession number NM_001134473.3) was cloned into the pLEX_307 (Addgene, #41392) and the pLEX_306 (Addgene, #41391) vector backbones, thus producing the pLEX_306 GSE1 and the pLEX_307 GSE1 constructs. *GSE1* coding sequence contained within the vector pENTR223 was first purchased by DNASU plasmid repository (HsCD00623069) ^55^: the plasmid was a kind gift from David Hill and David Root, from the Dana-Farber Cancer Institute, Broad Institute of Harvard and Massachusetts Institute of Technology (MIT), as a part of the ORFeome Collaboration ^56^. Subsequently, *GSE1* coding sequence was modified using the Q5 Site-directed Mutagenesis Kit (E0554S, New England Biolabs) to substitute the cytosine (C) in position 2495 with a thymine (T), a necessary step to generate the wild-type GSE1 transcript variant 2 (NM_001134473.3). At this point, the coding sequence was inserted through the Gateway System Technology into the pLEX_306 and pLEX_307 vectors ^57^.

### *In vivo* studies

*In vivo* studies were performed after approval from our animal facility and the institutional welfare committee “Organismo Preposto al Benessere degli Animali (OPBA)”. Experiments were notified to the Ministry of Health (as required by the Italian law; Institutional Animal Care and Use Committee numbers: 71/2019 in accordance with European Union directive 2010/63). NB4 cells were transduced with the empty pLKO.1 puro (EV) or pLKO.1 puro containing the shA1 and the shB2. After 24 hours, 1.5 x 10^6^ cells per mouse were re-suspended in 200 μl PBS containing 15% of Matrigel (Corning, 356231) and, then, injected subcutaneously in the left flank of 8-12 weeks old male and female NOD SCID IL2Rgnull (NSG) mice. For each mouse, the tumor size was measured three times per week with a linear caliper and the volume was calculated using the formula V = (a × b^2^)/2, where a and b are the longest and the shortest diameters of the tumor, respectively. Results are reported as tumor volume (mm^3^). Mice were sacrificed when the longest diameter of the tumor reached a size of approximately 15 mm.

### Flow cytometry analysis of cell surface markers

About 1 x 10^6^ cells were harvested, re-suspended in 300 μl of 5% BSA dissolved in PBS and blocked for 30 minutes at room temperature. Then, cells were re-pelleted and re-suspended in 100 μl primary antibody diluted in 1% BSA in PBS and let for 1h at room temperature in the dark. At this point, cells were washed with 1 ml of 1% BSA in PBS, centrifuged and re-suspended in 250 μl cold PBS. Transduced cells were, then, fixed by the addition of 250 μl 2% formaldehyde in PBS and incubated for 20 minutes in ice. After a further spinning, cell pellets were re-suspended in 500 μl PBS and analyzed by the FACS Celesta flow cytometer (BD Biosciences). Data analysis was performed using the FlowJo software. The antibodies used for the flow cytometry analysis of the cell surface markers were: anti-human CD11b (740965, BD OptiBuild™, 1:100), anti-human CD14 (11-0149-42, Thermo Fisher Scientific, 1:100), anti-human CD117 known also as cKIT (12-1178-42, Thermo Fisher Scientific, 1:100).

### Chromatin Immuno-Precipitation-sequencing (ChIP-seq)

ChIP-seq analysis was carried out in NB4 cells transduced with the pLEX_307 GSE1 and treated for 24 hours with either MC2580 or DMSO. As negative ChIP control we used NB4 cells transduced with the empty vector pLEX_307 (pLEX_307 EV). About 1 x 10^8^ cells for each condition were cross-linked by formaldehyde at 1% final concentration, which was added to the culture medium and incubated for 10 minutes with shaking. The reaction was stopped with 0.125 M Glycine and then samples were left shaking for 5 minutes at room temperature. Cells were then washed twice with PBS and lysed in SDS buffer (50 mM Tris-HCl pH 8.0, 0.5% SDS, 100 mM NaCl, 5 mM EDTA, 0.02% NaN3), supplemented with protease inhibitors (Roche, 04693116001) and 0.5 mM PMSF. At this point, we added the Triton Dilution buffer (100 mM NaCl, 100 mM Tris-HCl pH 8.5, 5 mM EDTA, 5% Triton X-100, 0.02% NaN3) supplemented with protease inhibitors (Roche, 04693116001) and 0.5 mM PMSF to the whole cell extracts to obtain the IP buffer conditions (100 mM NaCl, 33 mM Tris-HCl pH 8.0, 5 mM EDTA, 0.02% NaN3, 0.33% SDS, 1.7% Triton X-100, 33 mM Tris-HCl pH 8.5). Chromatin was then subjected to 35 cycles of sonication (30 seconds each) using a Branson Sonifier 250 to obtain DNA fragments of 300-bp average length. Subsequently, sheared chromatin was pre-cleared by incubation with 100 μl of Dynabeads protein G (Invitrogen, 10004D) for 2 hours on a rotating wheel at 4 °C. After preclearing, chromatin was used as input in the immuno-precipitation experiment, carried out overnight on a rotating wheel at 4 °C in the presence of 20 μg of anti-V5 antibody (Abcam, ab9116). 2.5% of the input sample was collected just before addition of the antibody and stored at −20 °C, for subsequent tests. After the overnight incubation, 200 μl of Dynabeads protein G were added to the antibody-chromatin reaction tube and incubated for 3 hours on a rotating wheel at 4 °C. Later, beads were washed thrice with Washing buffer A (1% Triton X-100, 0.1% SDS, 150mM NaCl, 2mM EDTA pH 8, 20mM Tris-HCl pH 8) and once with Washing buffer B (1% Triton X-100, 0.1% SDS, 500mM NaCl, 2mM EDTA pH 8, 20mM Tris-HCl pH 8) supplemented with protease inhibitors (Roche, 04693116001), followed by a final washing step with TE 1X buffer. At each wash, beads were incubated for 5 minutes on a rotating wheel at 4 °C. De-crosslinking/elution step of the ChIP samples were carried out by the addition of 300 μl de-crosslinking buffer (2% SDS, 0.25 mg/ml Proteinase K in TE 1X), followed by overnight incubation at 65 °C. The de-crosslinking step was also carried out for the input sample, by adding 3 volumes of de-crosslinking buffer to 1 volume of input. The day after, DNA from ChIPs and Input samples were purified using the DNA purification kit (QIAquick PCR Purification Kit, Qiagen, 28106) according to the manufacturer’s protocol. DNA libraries were prepared with 10 ng of DNA through an in-house protocol ^58^ by the IEO genomic facility and sequenced on a NovaSeq 6000 (Illumina) instrument.

### ChIP-seq data analysis

Short reads obtained from Illumina Genome Analyzer II were quality-filtered according to the ENCODE pipeline ^59^. Reads were aligned to the hg38 reference genome using Bowtie (v 4.8.2) ^60^. MACS (v 1.4.2) ^61^ was used as peakcaller to identify regions of ChIP-seq enrichment of the V5-GSE1 in DMSO and MC2580-treated cells, using as background both the respective input and the V5-ChIP performed in NB4 cells transduced with the pLEX_307 EV. Resulting enriched regions were annotated as specifically bound by GSE1 in each condition. Only reads with a unique match to the genome and with two or fewer mismatches (-m 1 –v 2) were retained. MACS p-value threshold was set to 10^−5^ for all the data sets. The four genomic region categories displayed in the Fig. 4D and the Fig. S10-S11 were obtained as follows: genomic regions bound by GSE1 in both DMSO- and MC2580-treated cells were defined as regions with peaks in both conditions and at least 1 bp of overlap between the two samples. These loci were further divided in “common same intensity”, that displayed a log2FC RPKM of DMSO vs MC2580 < 0.5 and “common MC2580 reduced intensity” showing a log2FC RPKM of DMSO vs MC2580 > 0.5. Regions commonly detected in both conditions and showing a log2FC RPKM of MC2580 vs DMSO > 0.5 were only 32 (“common DMSO reduced intensity”) and were not shown in the figures. “DMSO only” and “MC2580 only” categories instead contained genomic regions with identified peaks in DMSO or MC2580-treated cells, respectively. Within these regions, read counts were calculated with bedtools suite and only regions with a log2FC RPKM of DMSO vs MC2580 > 1 for “DMSO only” and log2FC RPKM of MC2580 vs DMSO > 1 for “MC2580 only” were kept. Annotation of the genomic regions in the different conditions was achieved using the R package ChIPseeker ^62^ and ChIPpeakAnno ^63^. The bigwig files for UCSC browser visualization of genome profiles were normalized with the deepToos suite. ChIP-seq data of LSD1, H3K4me1, H3K4me2, H3K4me3 and H3K27ac used for comparative analysis with V5-GSE1 ChIP-seq are described and present in ^29^.

## Acknowledgements

We thank R. Noberini for critical reading of the manuscript, Antonello Mai’s research group at Sapienza University of Rome and the Experimental Therapeutic Unit at IEO/IFOM campus for providing the MC2580 and DDP-38003 compounds, respectively; R. Giambruno, M. Marzi, F. Nicassio, M. Mihailovich, F. Santoro, the Mass Spectrometry, Genomic and Flow Cytometry Units at IEO for technical support and members of the Bonaldi’s group for helpful discussions on the project.

## Funding

T.B. research activity is supported by grants from the Italian Association for Cancer Research (grant# 15741) and by EPIC-XS, project number 823839, funded by the Horizon 2020 programme of the European Union; L.N. was supported by a Fellowship of the European Institute of Oncology foundation (FIEO) and was a PhD student within the European School of Molecular Medicine (SEMM); E.M. is supported by a fellowship of the Italian Foundation for Cancer Research (FIRC) and is a PhD student within the European School of Molecular Medicine (SEMM).

## Data and code availability

The accession number for all the RNA-seq data and the V5-GSE1 ChIP-seq data reported in this paper is GEO database: GSE164560.

The accession number for the LSD1, H3K4me1, H3K4me2, H3K4me3 and H3K27ac ChIP-seq data reported in ^29^ is GEO database: GSE128530.

The accession number for the MS-proteomics data reported in ^29^ is ProteomeXchange database: PXD012954.

## Conflict of interest

The authors declare that they have no conflict of interest.

## Notes

### Competing Interest Statement

The authors have declared no competing interest.

## References

1 Shi Y, Lan F, Matson C, Mulligan P, Whetstine JR, Cole PA et al. Histone demethylation mediated by the nuclear amine oxidase homolog LSD1. Cell 2004. doi:10.1016/j.cell.2004.12.012.

2 Metzger E, Wissmann M, Yin N, Müller JM, Schneider R, Peters AHFM et al. LSD1 demethylates repressive histone marks to promote androgen-receptor-dependent transcription. Nature 2005. doi:10.1038/nature04020.

3 Amente S, Lania L, Majello B. The histone LSD1 demethylase in stemness and cancer transcription programs. Biochim. Biophys. Acta - Gene Regul. Mech. 2013. doi:10.1016/j.bbagrm.2013.05.002.

4 Wang Y, Zhang H, Chen Y, Sun Y, Yang F, Yu W et al. LSD1 Is a Subunit of the NuRD Complex and Targets the Metastasis Programs in Breast Cancer. Cell 2009. doi:10.1016/j.cell.2009.05.050.

5 Humphrey GW, Wang Y, Russanova VR, Hirai T, Qin J, Nakatani Y et al. Stable Histone Deacetylase Complexes Distinguished by the Presence of SANT Domain Proteins CoREST/kiaa0071 and Mta-L1. J Biol Chem 2001. doi:10.1074/jbc.M007372200.

6 Lee MG, Wynder C, Cooch N, Shiekhattar R. An essential role for CoREST in nucleosomal histone 3 lysine 4 demethylation. Nature 2005. doi:10.1038/nature04021.

7 Shi YJ, Matson C, Lan F, Iwase S, Baba T, Shi Y. Regulation of LSD1 histone demethylase activity by its associated factors. Mol Cell 2005. doi:10.1016/j.molcel.2005.08.027.

8 Kim SA, Zhu J, Yennawar N, Eek P, Tan S. Crystal Structure of the LSD1/CoREST Histone Demethylase Bound to Its Nucleosome Substrate. Mol Cell 2020. doi:10.1016/j.molcel.2020.04.019.

9 Perillo B, Ombra MN, Bertoni A, Cuozzo C, Sacchetti S, Sasso A et al. DNA oxidation as triggered by H3K9me2 demethylation drives estrogen-induced gene expression. Science (80-) 2008. doi:10.1126/science.1147674.

10 Metzger E, Imhof A, Patel D, Kahl P, Hoffmeyer K, Friedrichs N et al. Phosphorylation of histone H3T6 by PKCB i controls demethylation at histone H3K4. Nature 2010. doi:10.1038/nature08839.

11 Nair SS, Nair BC, Cortez V, Chakravarty D, Metzger E, Schüle R et al. PELP1 is a reader of histone H3 methylation that facilitates oestrogen receptor-α target gene activation by regulating lysine demethylase 1 specificity. EMBO Rep 2010. doi:10.1038/embor.2010.62.

12 Schulte JH, Lim S, Schramm A, Friedrichs N, Koster J, Versteeg R et al. Lysine-specific demethylase 1 is strongly expressed in poorly differentiated neuroblastoma: Implications for therapy. Cancer Res 2009. doi:10.1158/0008-5472.CAN-08-1735.

13 Zhao ZK, Dong P, Gu J, Chen L, Zhuang M, Lu WJ et al. Overexpression of LSD1 in hepatocellular carcinoma: A latent target for the diagnosis and therapy of hepatoma. Tumor Biol 2013. doi:10.1007/s13277-012-0525-x.

14 Hayami S, Kelly JD, Cho HS, Yoshimatsu M, Unoki M, Tsunoda T et al. Overexpression of LSD1 contributes to human carcinogenesis through chromatin regulation in various cancers. Int J Cancer 2011. doi:10.1002/ijc.25349.

15 Lim S, Janzer A, Becker A, Zimmer A, Schüle R, Buettner R et al. Lysine-specific demethylase 1 (LSD1) is highly expressed in ER-negative breast cancers and a biomarker predicting aggressive biology. Carcinogenesis 2010. doi:10.1093/carcin/bgp324.

16 Harris WJ, Huang X, Lynch JT, Spencer GJ, Hitchin JR, Li Y et al. The Histone Demethylase KDM1A Sustains the Oncogenic Potential of MLL-AF9 Leukemia Stem Cells. Cancer Cell 2012. doi:10.1016/j.ccr.2012.03.014.

17 Lokken AA, Zeleznik-Le NJ. Breaking the LSD1/KDM1A Addiction: Therapeutic Targeting of the Epigenetic Modifier in AML. Cancer Cell. 2012. doi:10.1016/j.ccr.2012.03.027.

18 Schenk T, Chen WC, Göllner S, Howell L, Jin L, Hebestreit K et al. Inhibition of the LSD1 (KDM1A) demethylase reactivates the all-trans-retinoic acid differentiation pathway in acute myeloid leukemia. Nat Med 2012. doi:10.1038/nm.2661.

19 Maiques-Diaz A, Spencer GJ, Lynch JT, Ciceri F, Williams EL, Amaral FMR et al. Enhancer Activation by Pharmacologic Displacement of LSD1 from GFI1 Induces Differentiation in Acute Myeloid Leukemia. Cell Rep 2018. doi:10.1016/j.celrep.2018.03.012.

20 Fang J, Ying H, Mao T, Fang Y, Lu Y, Wang H et al. Upregulation of CD11b and CD86 through LSD1 inhibition promotes myeloid differentiation and suppresses cell proliferation in human monocytic leukemia cells. Oncotarget 2017. doi:10.18632/oncotarget.18564.

21 Chen Y, Jie W, Yan W, Zhou K, Xiao Y. Lysine-specific histone demethylase 1 (LSD1): A potential molecular target for tumor therapy. Crit Rev Eukaryot Gene Expr 2012. doi:10.1615/CritRevEukarGeneExpr.v22.i1.40.

22 Duan Y-C, Ma Y-C, Qin W-P, Ding L-N, Zheng Y-C, Zhu Y-L et al. Design and synthesis of tranylcypromine derivatives as novel LSD1/HDACs dual inhibitors for cancer treatment. Eur J Med Chem 2017. doi:10.1016/j.ejmech.2017.09.038.

23 Lillico R, Stesco N, Khorshid Amhad T, Cortes C, Namaka MP, Lakowski TM. Inhibitors of enzymes catalyzing modifications to histone lysine residues: Structure, function and activity. Future Med. Chem. 2016. doi:10.4155/fmc-2016-0021.

24 Zheng YC, Yu B, Chen ZS, Liu Y, Liu HM. TCPs: privileged scaffolds for identifying potent LSD1 inhibitors for cancer therapy. Epigenomics. 2016. doi:10.2217/epi-2015-0002.

25 Lee MG, Wynder C, Schmidt DM, McCafferty DG, Shiekhattar R. Histone H3 Lysine 4 Demethylation Is a Target of Nonselective Antidepressive Medications. Chem Biol 2006. doi:10.1016/j.chembiol.2006.05.004.

26 Binda C, Valente S, Romanenghi M, Pilotto S, Cirilli R, Karytinos A et al. Biochemical, structural, and biological evaluation of tranylcypromine derivatives as inhibitors of histone demethylases LSD1 and LSD2. J Am Chem Soc 2010. doi:10.1021/ja101557k.

27 Vianello P, Botrugno OA, Cappa A, Dal Zuffo R, Dessanti P, Mai A et al. Discovery of a Novel Inhibitor of Histone Lysine-Specific Demethylase 1A (KDM1A/LSD1) as Orally Active Antitumor Agent. J Med Chem 2016; 59: 1501–1517.

28 Zheng YC, Ma J, Wang Z, Li J, Jiang B, Zhou W et al. A Systematic Review of Histone Lysine-Specific Demethylase 1 and Its Inhibitors. Med Res Rev 2015. doi:10.1002/med.21350.

29 Ravasio R, Ceccacci E, Nicosia L, Hosseini A, Rossi PL, Barozzi I et al. Targeting the scaffolding role of LSD1 (KDM1A) poises acute myeloid leukemia cells for retinoic acid-induced differentiation. Sci Adv 2020. doi:10.1126/sciadv.aax2746.

30 Ishikawa Y, Gamo K, Yabuki M, Takagi S, Toyoshima K, Nakayama K et al. A Novel LSD1 Inhibitor T-3775440 Disrupts GFI1B-Containing Complex Leading to Transdifferentiation and Impaired Growth of AML Cells. Mol Cancer Ther 2017. doi:10.1158/1535-7163.MCT-16-0471.

31 Chai P, Tian J, Zhao D, Zhang H, Cui J, Ding K et al. GSE1 negative regulation by miR-489-5p promotes breast cancer cell proliferation and invasion. Biochem Biophys Res Commun 2016. doi:10.1016/j.bbrc.2016.01.168.

32 Ding K, Tan S, Huang X, Wang X, Li X, Fan R et al. GSE1 predicts poor survival outcome in gastric cancer patients by SLC7A5 enhancement of tumor growth and metastasis. J Biol Chem 2018. doi:10.1074/jbc.RA117.001103.

33 Huang M, Tailor J, Zhen Q, Gillmor AH, Miller ML, Weishaupt H et al. Engineering Genetic Predisposition in Human Neuroepithelial Stem Cells Recapitulates Medulloblastoma Tumorigenesis. Cell Stem Cell 2019; 25: 433–446.e7.

34 Sehrawat A, Gao L, Wang Y, Bankhead A, McWeeney SK, King CJ et al. LSD1 activates a lethal prostate cancer gene network independently of its demethylase function. Proc Natl Acad Sci 2018. doi:10.1073/pnas.1719168115.

35 Hakimi MA, Dong Y, Lane WS, Speicher DW, Shiekhattar R. A candidate X-linked mental retardation gene is a component of a new family of histone deacetylase-containing complexes. J Biol Chem 2003. doi:10.1074/jbc.M208992200.

36 Yamamoto R, Kawahara M, Ito S, Satoh J, Tatsumi G, Hishizawa M et al. Selective dissociation between LSD1 and GFI1B by a LSD1 inhibitor NCD38 induces the activation of ERG super-enhancer in erythroleukemia cells. Oncotarget 2018. doi:10.18632/oncotarget.24774.

37 McClellan D, Casey MJ, Bareyan D, Lucente H, Ours C, Velinder M et al. Growth Factor Independence 1B-Mediated Transcriptional Repression and Lineage Allocation Require Lysine-Specific Demethylase 1-Dependent Recruitment of the BHC Complex. Mol Cell Biol 2019. doi:10.1128/mcb.00020-19.

38 Zhang X, Smits AH, Van Tilburg GBA, Ovaa H, Huber W, Vermeulen M. Proteome-wide identification of ubiquitin interactions using UbIA-MS. Nat Protoc 2018. doi:10.1038/nprot.2017.147.

39 Perillo B, Tramontano A, Pezone A, Migliaccio A. LSD1: more than demethylation of histone lysine residues. Exp Mol Med 2020. doi:10.1038/s12276-020-00542-2.

40 Carnesecchi J, Cerutti C, Vanacker JM, Forcet C. ERRα protein is stabilized by LSD1 in a demethylation-independent manner. PLoS One 2017. doi:10.1371/journal.pone.0188871.

41 Chao A, Lin CY, Chao AN, Tsai CL, Chen MY, Lee LY et al. Lysine-specific demethylase 1 (LSD1) destabilizes p62 and inhibits autophagy in gynecologic malignancies. Oncotarget 2017. doi:10.18632/oncotarget.20158.

42 Lan H, Tan M, Zhang Q, Yang F, Wang S, Li H et al. LSD1 destabilizes FBXW7 and abrogates FBXW7 functions independent of its demethylase activity. Proc Natl Acad Sci U S A 2019. doi:10.1073/pnas.1902012116.

43 Mihailovich M, Bremang M, Spadotto V, Musiani D, Vitale E, Varano G et al. MiR-17-92 fine-tunes MYC expression and function to ensure optimal B cell lymphoma growth. Nat Commun 2015. doi:10.1038/ncomms9725.

44 Supek F, Bošnjak M, Škunca N, Šmuc T. Revigo summarizes and visualizes long lists of gene ontology terms. PLoS One 2011. doi:10.1371/journal.pone.0021800.

45 Croft D, O’Kelly G, Wu G, Haw R, Gillespie M, Matthews L et al. Reactome: A database of reactions, pathways and biological processes. Nucleic Acids Res 2011. doi:10.1093/nar/gkq1018.

46 Zhang X, Tan Z, Kang T, Zhu C, Chen S. Arsenic sulfide induces miR-4665-3p to inhibit gastric cancer cell invasion and migration. Drug Des Devel Ther 2019. doi:10.2147/DDDT.S209219.

47 Sonenberg N, Hinnebusch AG. Regulation of Translation Initiation in Eukaryotes: Mechanisms and Biological Targets. Cell. 2009. doi:10.1016/j.cell.2009.01.042.

48 Li Y, Tao L, Zuo Z, Zhou Y, Qian X, Lin Y et al. ZY0511, a novel, potent and selective LSD1 inhibitor, exhibits anticancer activity against solid tumors via the DDIT4/mTOR pathway. Cancer Lett 2019. doi:10.1016/j.canlet.2019.03.052.

49 Abdel-Aziz AK, Pallavicini I, Ceccacci E, Meroni G, Saadeldin MK, Varasi M et al. Tuning mTORC1 activity dictates the response to LSD1 inhibition of acute myeloid leukemia. Haematologica 2020. doi:10.3324/haematol.2019.224501.

50 Wang X, Proud CG. The mTOR pathway in the control of protein synthesis. Physiology. 2006. doi:10.1152/physiol.00024.2006.

51 Dobin A, Davis CA, Schlesinger F, Drenkow J, Zaleski C, Jha S et al. STAR: Ultrafast universal RNA-seq aligner. Bioinformatics 2013. doi:10.1093/bioinformatics/bts635.

52 Li B, Dewey CN. RSEM: Accurate transcript quantification from RNA-Seq data with or without a reference genome. BMC Bioinformatics 2011. doi:10.1186/1471-2105-12-323.

53 Love MI, Huber W, Anders S. Moderated estimation of fold change and dispersion for RNA-seq data with DESeq2. Genome Biol 2014. doi:10.1186/s13059-014-0550-8.

54 Chen EY, Tan CM, Kou Y, Duan Q, Wang Z, Meirelles G V. et al. Enrichr: Interactive and collaborative HTML5 gene list enrichment analysis tool. BMC Bioinformatics 2013. doi:10.1186/1471-2105-14-128.

55 Seiler CY, Park JG, Sharma A, Hunter P, Surapaneni P, Sedillo C et al. DNASU plasmid and PSI:Biology-Materials repositories: Resources to accelerate biological research. Nucleic Acids Res 2014. doi:10.1093/nar/gkt1060.

56 Wiemann S, Pennacchio C, Hu Y, Hunter P, Harbers M, Amiet A et al. The ORFeome Collaboration: A genome-scale human ORF-clone resource. Nat. Methods. 2016. doi:10.1038/nmeth.3776.

57 Liang X, Peng L, Baek CH, Katzen F. Single step BP/LR combined Gateway reactions. Biotechniques 2013. doi:10.2144/000114101.

58 Blecher-Gonen R, Barnett-Itzhaki Z, Jaitin D, Amann-Zalcenstein D, Lara-Astiaso D, Amit I. High-throughput chromatin immunoprecipitation for genome-wide mapping of in vivo protein-DNA interactions and epigenomic states. Nat Protoc 2013. doi:10.1038/nprot.2013.023.

59 Landt SG, Marinov GK, Kundaje A, Kheradpour P, Pauli F, Batzoglou S et al. ChIP-seq guidelines and practices of the ENCODE and modENCODE consortia. Genome Res. 2012. doi:10.1101/gr.136184.111.

60 Langmead B, Trapnell C, Pop M, Salzberg SL. Ultrafast and memory-efficient alignment of short DNA sequences to the human genome. Genome Biol 2009. doi:10.1186/gb-2009-10-3-r25.

61 Zhang Y, Liu T, Meyer CA, Eeckhoute J, Johnson DS, Bernstein BE et al. Model-based analysis of ChIP-Seq (MACS). Genome Biol 2008. doi:10.1186/gb-2008-9-9-r137.

62 Yu G, Wang LG, He QY. ChIP seeker: An R/Bioconductor package for ChIP peak annotation, comparison and visualization. Bioinformatics 2015. doi:10.1093/bioinformatics/btv145.

63 Zhu LJ, Gazin C, Lawson ND, Pagès H, Lin SM, Lapointe DS et al. ChIPpeakAnno: A Bioconductor package to annotate ChIP-seq and ChIP-chip data. BMC Bioinformatics 2010. doi:10.1186/1471-2105-11-237.

64 Ritchie ME, Phipson B, Wu D, Hu Y, Law CW, Shi W et al. Limma powers differential expression analyses for RNA-sequencing and microarray studies. Nucleic Acids Res 2015. doi:10.1093/nar/gkv007.

