## Supplementary Materials and Methods and Figures for "Pharmacological inhibition of LSD1 triggers myeloid differentiation by targeting GSE1, a novel oncogene in AML"

The PDF file contains: Supplementary Materials and Methods, Supplementary Figures from S1 to S14 with the corresponding legends, legends of the Table S1, S2, S3, S4, S5 and S6 (these tables are also included as Supplementary Materials), References.

### Supplementary Materials and Methods

#### Protein co-immunoprecipitation (co-IP)

Subcellular fractionation, followed by protein co-immunoprecipitation (co-IP) using the nuclear extract as input were performed as already described in [1]. The antibodies employed for the co-IPs using LSD1 and GSE1 as baits were anti-LSD1 (Abcam, ab17721) and anti-GSE1 (Proteintech, 24947-1-AP), respectively.

**Differential Enrichment Analysis of Proteomics Data (DEP) of LSD1 interactome after LSD1 pharmacological inhibition**

The LSD1 interactome dataset upon 24 hours treatment with MC2580 was published in [1] and here it was re-analysed using DEP [2]. For this analysis, the intensity in the heavy and light channels of the two SILAC replicates (Forward and Reverse) was used to calculate protein changes. A protein was considered significantly enriched/depleted if its $\log_2$ fold change was greater than 1.5 or lower than -1.5 and its p-value (corrected for multiple testing) was < 0.05.

**Western blot analysis**

Cells were harvested, washed twice with PBS and re-suspended in RIPA buffer (10 mM Tris pH 8, 150 mM NaCl, 0.1% SDS, 1% Triton X-100, 1 mM EDTA, 0.1% Sodium Deoxicholate), supplemented with 1X EDTA-free Protease Inhibitors (Roche, 11836170001), 0.5 mM PMSF, 1 mM DTT and 1:50 Benzonase (250U/µl, E1014, Millipore). Re-suspended cells were incubated 45 minutes at room temperature on a rotating wheel and vortexed briefly at 15-minute intervals to facilitate cell lysis and protein extraction. Afterwards, samples were spun down at 16 060 g for 30 minutes, at 4 °C in a bench-top centrifuge and supernatants -corresponding to the whole protein extracts- were collected and quantified by BCA assay (23225, Thermo Fisher Scientific). Around 30-40 µg of protein extract was mixed with Laemmli buffer containing 100 mM DTT and denatured for 5 minutes at 95°C before loading on SDS-PAGE gel. Transfer of protein to PVDF membrane was performed at 100 V for 1 hour and 30 minutes at 4°C, or at 30 V overnight at 4°C in Transfer buffer (Tris-Glycine) containing 10% methanol. Membranes were blocked in 10% BSA dissolved in TBS/0.1% Tween for 2 hours at room temperature. Primary antibodies were diluted in 5% BSA in TBS/0.1% Tween and incubated with the PVDF membrane overnight at 4°C, or for 3 hours at room temperature. After three washes with TBS/0.1% Tween (5 minutes each), membranes were incubated with the corresponding secondary antibody in 5% BSA in TBS/0.1% Tween for 1 hour at room temperature. After three additional washes, protein signals were detected using the Enhanced Chemo Luminescence (ECL) method (Biorad, 1705061) and images were acquired with the ChemiDoc XRS+ System (Biorad, 1708265). Quantitation of the bands was carried out with the ImageJ software. The following antibodies were used according to the company's instruction at the indicated dilutions: anti-LSD1 (Abcam, ab17721, 1:1000), anti-GSE1 (Proteintech, 24947-1-AP, 1:600), anti-GFI1 (Santa Cruz Technology, sc-376949, 1:100), anti-H3 (Abcam, ab1791, 1:5000), anti-Vinculin (Sigma Aldrich, V9131, 1:5000), anti c-MYC (Santa Cruz Technology, sc-764, 1:200), anti-cleaved caspase 3 (Cell Signalling Technology, #9661S, 1:1000), anti-total caspase 3 (Cell Signalling Technology, #9662S, 1:1000), anti-V5 (Invitrogen, R960-25, 1:5000).

**RNA extraction and RT-qPCR**

Total RNA was extracted by using the RNeasy Mini kit (Qiagen, 74104) and reverse transcription was performed with SuperScript™ II Reverse Transcriptase (Invitrogen, 18064014), according to the manufacturer's protocols. Real time PCR was carried out in triplicate, in 20 μl final reaction volume constituted by 10 μl of 2X SYBR green Master Mix (Applied Biosystems, 4385614), 20 ng of cDNA retro-transcribed from the RNA, and 0.5 μM of each primer mix. Quantitative PCR amplifications were performed in the CFX96 Real-Time System (BioRad), using the following protocol: 1) 95 °C for 2 minutes, 2) 40 cycles at 95 °C for 10 seconds and 60 °C for 30 seconds. The sequences of the primers are listed in the Table S6.

**Cell Transduction**

Lentiviral constructs were transiently transfected in HEK-293T cells by calcium phosphate transfection method [3] using the packaging plasmid pCMV-DR8.74 and the envelope plasmid pMD2G-VSVG. The retroviral vectors were, instead, transiently transfected with the same method in phoenix-AMPHO cells, together with the packaging plasmid pKAT. Supernatants from the HEK-293T or Phoenix-AMPHO cells were filtered with 0.45 μm filter and then ultra-centrifuged at 70 770 x g for 2 hours at 4 °C. The pellet -enriched of viral particles- was resuspended in culture medium and added to 2 x $10^6$ NB4 or THP-1 cells, previously plated in 6-well plates at 1 x $10^6$/ml concentration, together with 16 μg of polybrene. Cells were centrifuged at 1000 x g for 1 hour at room temperature in a swing-rotor centrifuge (Spin Infection) and then left incubating for 4 hours at 37 °C, prior to the addition of 2 ml growth medium. In the meantime, transfected HEK-293T or Phoenix-AMPHO cells were replaced with fresh medium that was, again, filtered and ultra-centrifuged for the second round of infection carried out the day after following the same protocol. Lentiviral and retroviral transductions were carried out in the Biosafety level 2 laboratory (BSL-2).

**Cell growth and viability assays**

For the growth curve, cultured cells were counted in Trypan Blue solution (T8154, Sigma Aldrich) using the TC20 automated cell counter from Bio-Rad (http://www.bio-rad.com/it-it/product/tc20-automated-cell-counter). The percentage of living cells was also assessed with this method.

**Cell cycle analysis by flow cytometry**

Cells were harvested, washed once with 1% BSA in PBS and re-suspended in 250 μl PBS. Cell fixing was carried out by adding 750 μl of pure ethanol dropwise, upon vortexing. After 30 minutes, cells were washed again with 1% BSA in PBS and re-suspended in Propidium Iodide (PI, 20 μg/ml) + RNase A (250 μg/ml). Stained cells were let on fluorescence activated cell sorting (FACS) tubes overnight at 4 °C before analysing the samples with the FACS Celesta flow cytometer (BD Biosciences). Analysis of the cell cycle was performed using the function “cell cycle” of the FlowJo software.

**ChIP-qPCR**

For validation of specific genomic loci extrapolated from the ChIP-seq data, ChIP-qPCR was performed as follows: 1 µl of DNA from ChIP and input samples was mixed with 10 µl of 2X SYBR green Master Mix (Applied Biosystems, 4385614) and 0.5 µM of each primer mix in a final reaction volume of 20 µl. As previously described, real time PCR was performed in three technical replicates and quantitative PCR amplifications were carried out in the CFX96 Real-Time System (BioRad), using the following conditions: 1) 95 °C for 2 minutes, 2) 40 cycles at 95 °C for 10 seconds and 60 °C for 30 seconds. The primers used are listed in the Table S6.

**Statistical analysis**

Most of the data are represented as mean + standard deviation (SD) and statistical analyses were carried out using two-tailed paired Student's t-tests or one sample t-tests, unless others specified. The number (*n*) of biological replicates, the type of statistical analyses performed, and statistical significance are reported in the corresponding figures and/or figure legends for each experiment.

### Supplementary Figures

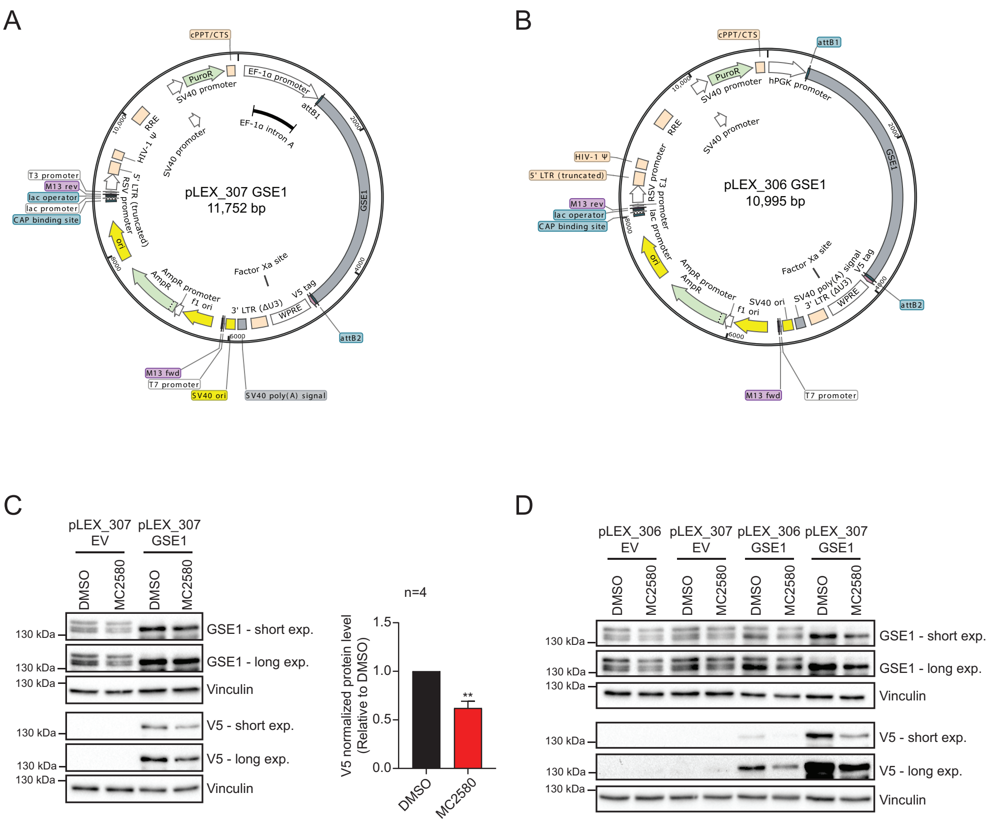


**Fig. S1: MC2580 inhibitor decreases the protein expression of an exogenous form of GSE1.** A) Map of the pLEX_307 vector in which the insert of the GSE1 coding sequence (NM_001134473.3) tagged with the V5 at the C-terminus, has been cloned. B) Map of the pLEX_306 vector in which the insert of GSE1 coding sequence (NM_001134473.3) tagged with the V5 at the C-terminus, has been cloned. C) *Left panel:* Western Blot analysis of GSE1 and V5 in NB4 cells transduced with either the empty pLEX_307 (pLEX_307 EV), or the pLEX_307 GSE1 and treated for 24 hours with MC2580 (2 µM) and DMSO. Vinculin was used as loading control. *Right panel:* Bar-graph displaying quantitation of the exogenous V5-tagged GSE1 protein level, normalized over the Vinculin, in 4 independent replicates of pLEX_307 EV and pLEX_307 GSE1 transduced NB4 cells treated for 24 hours with either MC2580 (2 µM) or DMSO. The results are plotted as fold change of V5-GSE1 protein level in the treated sample normalised over the level of the corresponding protein in the control, DMSO-treated cells. The chart represents mean + standard deviation from n=4 biological replicates (one sample T-test, **p value<0.01). D) Western Blot analysis of GSE1 and V5 in pLEX_306 EV, pLEX_307 EV, pLEX_306 GSE1 and pLEX_307 GSE1 transduced in NB4 cells and treated for 24 hours with MC2580 (2 μM) and DMSO as negative control. Vinculin was used as loading control.

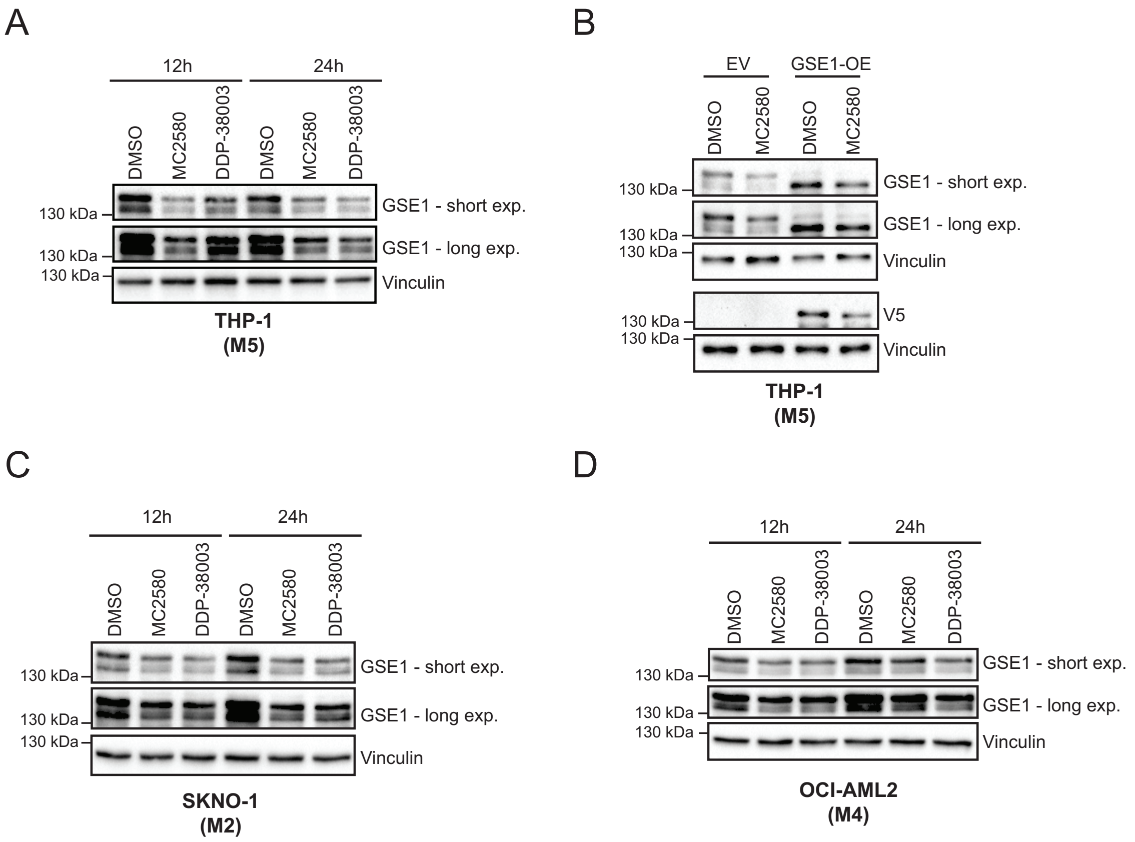


**Fig. S2: LSD1 inhibitors reduce GSE1 protein level in other AML subtypes beyond APL.** A) Western Blot analysis of GSE1 in THP-1 cells treated with MC2580 (2 µM), DDP-38003 (2 µM) and DMSO for 12 and 24 hours. Vinculin was used as loading control. M5 refers to the AML subtype to which these cells belong, according to the French-American-British (FAB) classification. B) Western Blot analysis of GSE1 and V5 in control (pLEX_307 EV) and GSE1 over-expressing THP-1 cells generated by lentiviral transduction of the pLEX_307 GSE1. Cells were subsequently treated with MC2580 (2 µM) or DMSO for 24 hours. Vinculin was used as loading control. M5 refers to the AML subtype in which these cells are included, according to the FAB classification. C) Western Blot analysis of GSE1 in SKNO-1 cells treated with MC2580 (2 µM), DDP-38003 (2 µM) and DMSO for 12 and 24 hours. Vinculin was used as loading control. M2 refers to the AML subtype in which these cells are included, according to the FAB classification. D) Western Blot analysis of GSE1 in OCI-AML2 cells treated with MC2580 (2 µM), DDP-38003 (2 µM) and DMSO for 12 and 24 hours. Vinculin was used as loading control. M4 refers to the AML subtype to which these cells belong, according to the FAB classification.

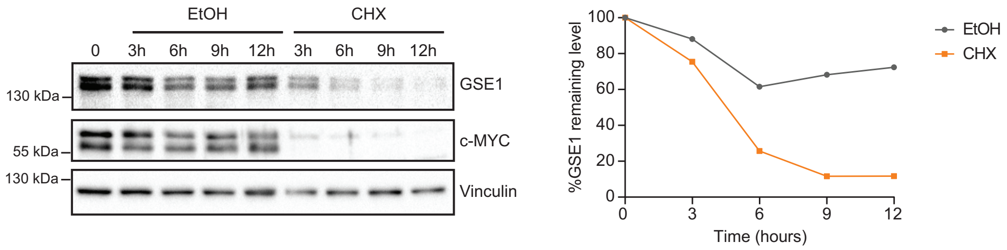


**Fig. S3: Analysis of GSE1 protein stability in NB4 cells.** *Left panel:* Time-resolved Western Blot analysis of GSE1 and c-MYC protein levels in NB4 cells at 3-, 6-, 9- and 12-hours post-treatment with cycloheximide (CHX) (0.1 mg/ml) and Ethanol (EtOH) as negative control. Vinculin was used as loading control. *Right panel:* Line-plot displaying the percentage of GSE1 protein level normalized over Vinculin in cycloheximide-treated NB4 cells, compared to control, EtOH-treated ones.

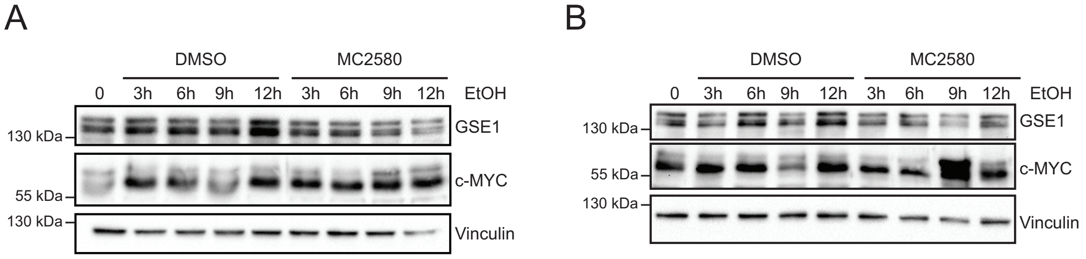


**Fig. S4: Assessment of the efficiency of MC2580 treatment in the absence of cycloheximide as control for the results displayed in figure 1H.** A) Time-resolved Western Blot analysis of GSE1 and c-MYC levels in NB4 cells treated with EtOH (negative control for CHX), in combination with either MC2580 (2 µM) or DMSO for 3-, 6-, 9- and 12-hours as control of the first biological replicate of the experiment displayed in Figure 1H. Vinculin was used as loading control. B) Western Blot analysis of GSE1 and c-MYC levels in NB4 cells treated with EtOH in combination with either MC2580 (2 µM) or DMSO for 3, 6, 9 and 12 hours as control of the second biological replicate of the experiment in Figure 1H. Vinculin was used as loading control.

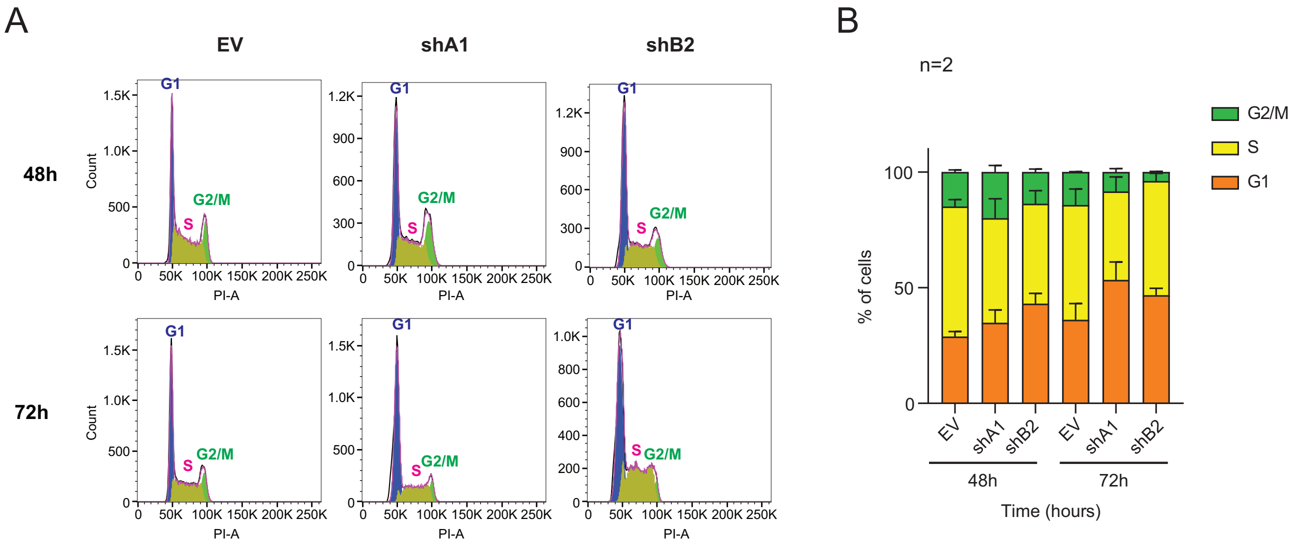


**Fig. S5: GSE1 knock down (KD) increases the percentage of cells in the G1-phase.** A) Representative cell cycle flow cytometry analysis of NB4 cells at 48- and 72-hours post-transduction with either an empty pLKO.1 puro vector (EV) or the pLKO.1 puro vector, in which the shA1 and shB2 constructs targeting GSE1 are cloned. B) Percentage of shA1, shB2 and EV-transduced cells in each cell cycle phase at 48 and 72 hours upon transduction. The chart represents mean + standard deviation (SD) of two biological replicates (n=2).

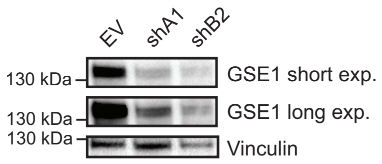


**Fig. S6: Western Blot validation of GSE1 KD in cells used for transplantation in NOD/SCID gamma (NSG) mice.** Western Blot analysis of GSE1 in NB4 cells 48- hours post transduction with either the empty pLKO.1 puro vector (EV) or the same vector containing the shA1 and shB2 constructs targeting GSE1. Vinculin was used as loading control.

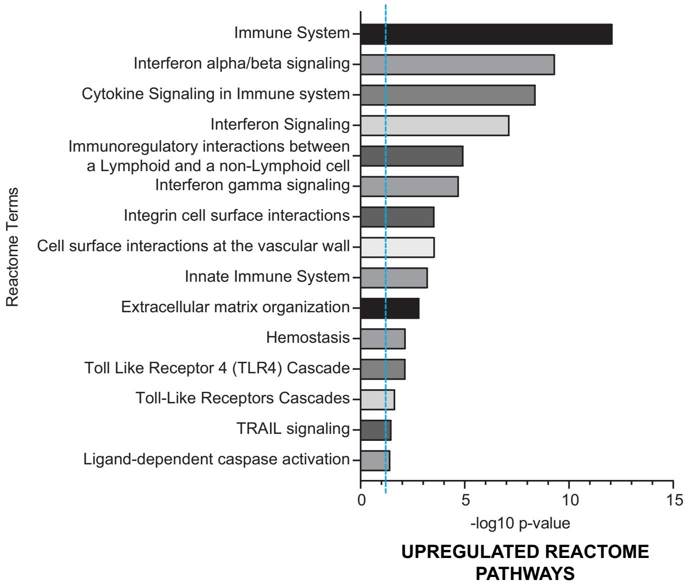


**Fig. S7: Reactome analysis of the 422 up-regulated genes upon GSE1 KD.** Bar-graph displaying the Reactome results on the enriched pathways associated with the 422 up-regulated genes upon GSE1 depletion in NB4 cells. Reactome analysis and calculation of the statistically significant pathway terms was performed with EnrichR [4] (Adjusted p-value < 0.05). The x-axis displays the -$\log_{10}$ of the adjusted p-value for each significant term. The blue dashed line indicates the significant threshold used to define the enriched pathways.

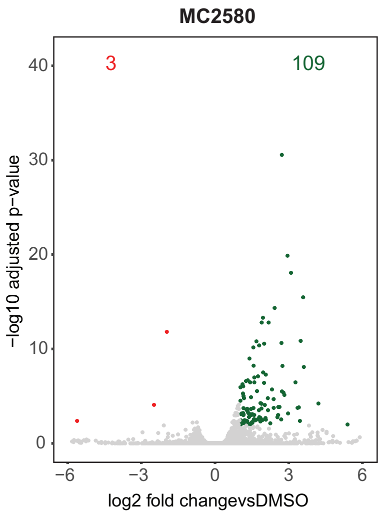


**Fig. S8: Analysis of differentially expressed genes upon MC2580 treatment in NB4 cells.** Volcano-plot displaying the up- and down- regulated genes 24 hours post- MC2580 treatment. On the x- axis, the $\log_2$ fold change (FC) values of LSD1-inhibited cells versus DMSO-treated ones are displayed, while y-axis displays the -$\log_{10}$ of the adjusted p-value (padj) for each gene. FC value was calculated with DEseq2 program, using two biological replicates for each condition (n=2).

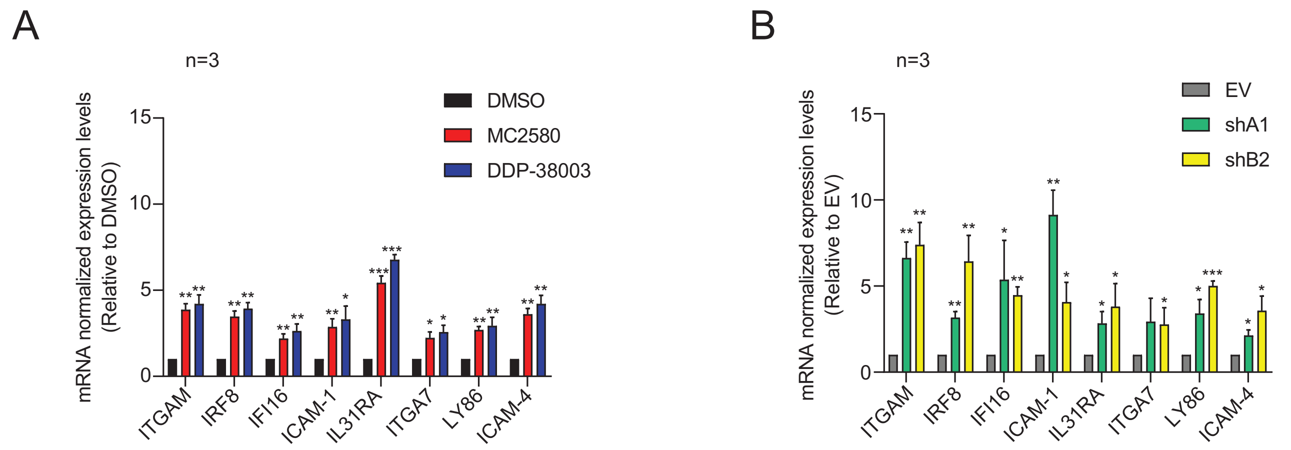


**Fig. S9: RT-qPCR analysis validates RNA-seq results.** A) RT-qPCR analysis on a panel of genes in NB4 cells treated for 24 hours with MC2580 (2 µM), DDP-38003 (2 µM) and DMSO as control. Ct values of different genes are normalized against GAPDH. The results are plotted as fold change (FC) of the mRNA levels in the treated conditions compared to DMSO. Chart represents mean + standard deviation (SD) (n=3 biological replicates, one sample T-test, ***p-value<0.001, **p-value<0.01, *p-value<0.05). B) RT-qPCR analysis on a panel of genes at 48 hours post-transduction with shA1, shB2 and the empty vector pLKO.1 puro (EV) as negative control. Ct values of the different genes are normalized over GAPDH. The results are plotted as FC of the mRNA levels in the GSE1 KD samples over the corresponding levels in the EV-transduced cells. Bar-graph represents mean + SD (n=3 biological replicates, one sample T-test, ***p-value<0.001, **p-value<0.01, *p-value<0.05).

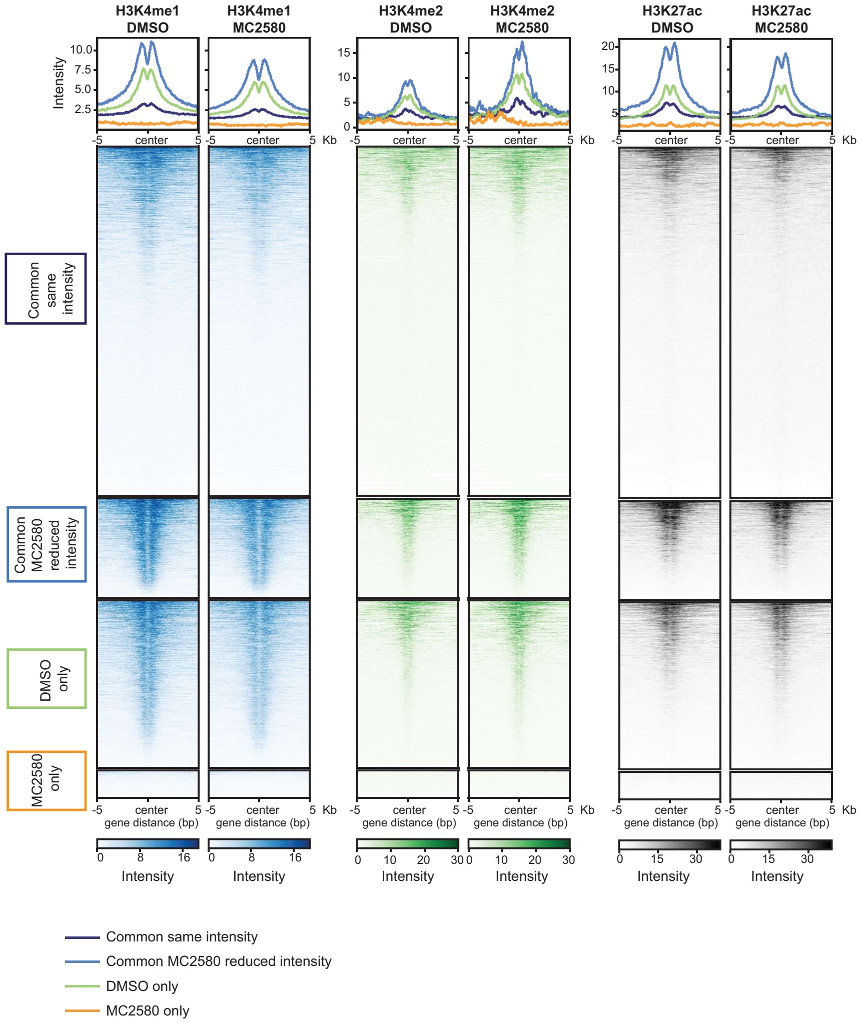


**Fig. S10: Effect of MC2580 treatment on H3K4me1, H3K4me2 and H3K27ac localisation at the GSE1-bound genomic loci.** Heatmap representation of the normalized ChIP-seq intensities of H3K4me1, H3K4me2 and H3K27ac at ±5 kb around the center of the GSE1-bound loci, 24 hours after treatment with MC2580 (2 μM) and DMSO. The heatmap is divided in different groups, as classified and reported in Figure 4D. The H3K4me1, H3K4me2 and H3K27ac ChIP-seq data used has already been published in [1].

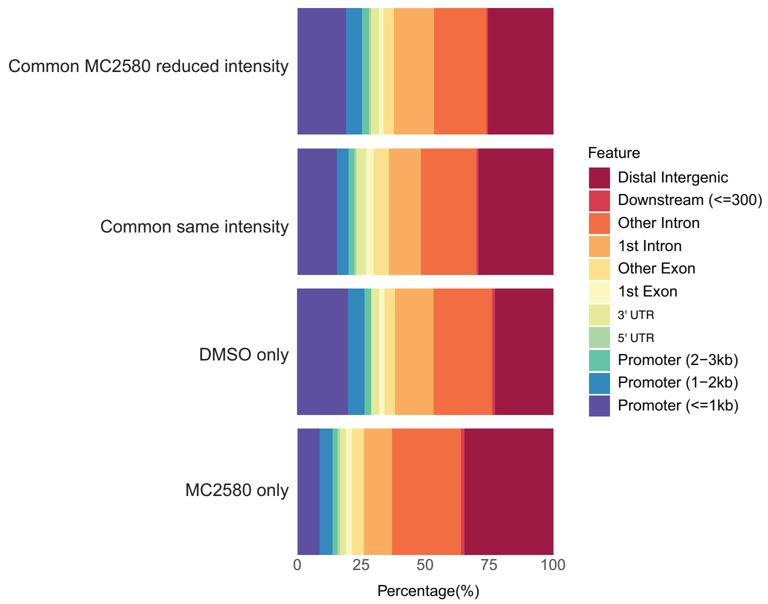


**Fig. S11: GSE1 binding at promoters is reduced in the "Common same intensity" and "MC2580 only" subsets.** Genome annotation of the V5-GSE1 ChIP-seq peaks in the different genomic categories described in Figure 4D.

***Physiological System Development and Function***

**MC2580**

| Name | p-value range | #Molecules |
|---|---|---|
| Hematological System Development and Function | 2.09E-03 - 3.96E-10 | 42 |
| Immune Cell Trafficking | 2.05E-03 - 7.86E-10 | 32 |
| Connective Tissue Development and Function | 2.06E-03 - 1.87E-09 | 21 |
| Skeletal and Muscular System Development and Function | 2.12E-03 - 1.87E-09 | 22 |
| Tissue Development | 2.12E-03 - 1.87E-09 | 48 |

**GSE1 KD**

| Name | p-value range | #Molecules |
|---|---|---|
| Hematological System Development and Function | 3.00E-07 - 6.25E-32 | 171 |
| Tissue Morphology | 2.91E-07 - 6.25E-32 | 158 |
| Immune Cell Trafficking | 3.00E-07 - 3.32E-29 | 123 |
| Lymphoid Tissue Structure and Development | 2.91E-07 - 1.66E-27 | 137 |
| Hematopoiesis | 2.25E-07 - 9.99E-23 | 80 |

**Fig. S12: Ingenuity Pathway Analysis (IPA) of "Physiological System Development and Function" terms upon LSD1 inhibition and GSE1 KD.** Representation of the top five "Physiological System Development and Function" IPA terms extracted from the analysis of the differentially expressed genes (DEGs) upon MC2580 treatment and GSE1 KD.

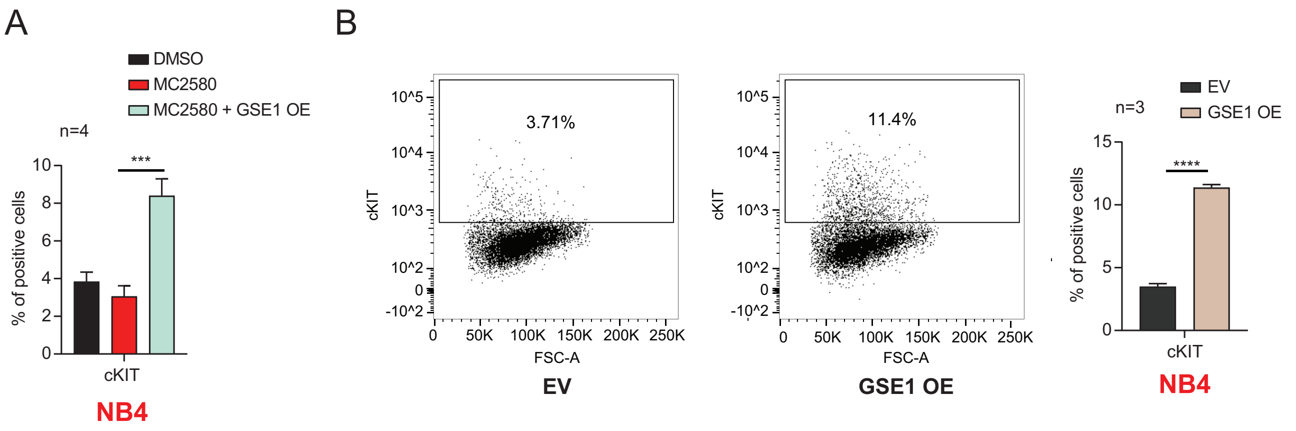


**Fig. S13: GSE1 over-expression (OE) in NB4 cells increases cKIT stemness marker.** A) Flow cytometry analysis to estimate the percentage of cKIT-positive cells in control (EV) and GSE1-overexpressing (OE) NB4 cells, treated for 24 hours with MC2580 (2 µM). GSE1 OE NB4 cells were generated by transducing the pLEX_307 GSE1, while control cells were transduced with the empty vector (EV). Chart represents mean + standard deviation (SD) (n=4 biological replicates; paired T-test, ***p-value<0.001). B) *Left panel:* The dot-plot displays the percentage of cKIT-positive cells in EV and GSE1 OE NB4 cells from one of the four biological replicates and used as representative result. *Right panel:* Bar-graph of the quantitation of the percentage of cKIT-positive cells in EV and GSE1 OE NB4 cells. Chart represents mean + standard deviation (SD) (n=3 biological replicates; unpaired T-test, ****p-value<0.0001).

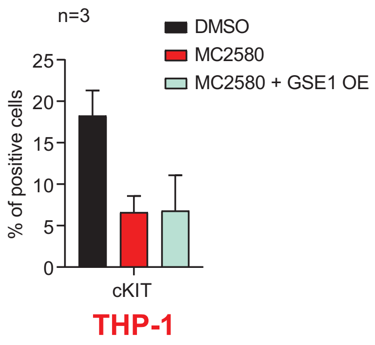


**Fig. S14: GSE1 OE does not rescue the down-regulation of cKIT triggered by MC2580 in THP-1 cell line.** Flow cytometry analysis of cKIT-positive cells in control (EV) and GSE1 OE THP-1 cells treated for 24 hours with MC2580 (2 μM). GSE1 OE THP-1 cells were produced by lentiviral transduction of the pLEX_307 GSE1, while control cells were transduced with the empty vector (EV). The chart represents mean + standard deviation (SD) (n=3 biological replicates).

**Supplementary Table Legends**

**Table S1:** Gene expression profiling by RNA-seq in NB4 cells transduced for 48 hours with shA1 and shB2 targeting GSE1 or the empty pLKO.1 lentiviral vector as negative control.

**Table S2:** Gene Ontology (GO) analysis of the Biological Processes (BP) enriched after GSE1 knock-down (KD). *Spreadsheet 1*: Complete list of BP terms obtained from the GO analysis of the up-regulated genes after GSE1 KD. Statistically significant terms are displayed in green (adjusted p-value < 0.05). GO analysis was performed through EnrichR [4]. *Spreadsheet 2*: List of BP terms extrapolated from the GO analysis of the down-regulated genes after GSE1 KD. GO analysis was performed through EnrichR [4].

**Table S3:** RNA-seq analysis of NB4 cells treated for 24 hours with 2 μM of MC2580 or DMSO.

**Table S4:** Ingenuity Pathway Analysis (IPA) of the differentially expressed genes (DEG) after LSD1 pharmacological inhibition and GSE1 knock-down (KD). *Spreadsheet 1*: List of the IPA pathways extrapolated from the analysis of the DEG after 24h treatment with MC2580 (2 μM). Statistically significant IPA pathways are displayed in green ($-\log_{10}$ p-value > 2). Associated biological processes to each IPA term are annotated manually and indicated only for the significant IPA pathways. *Spreadsheet 2*: List of the IPA pathways obtained from the analysis of the DEG upon GSE1 KD. Statistically significant IPA pathways are displayed in green ($-\log_{10}$ p-value > 2). Associated biological processes to each IPA term are annotated manually and indicated only for the significant IPA pathways.

**Table S5:** Annotation of the V5-GSE1 binding sites in the genomic categories "Regions bound by GSE1 similarly in both DMSO- and MC2580- treated cells" (*Spreadsheet 1*), "Regions bound by GSE1 only in DMSO condition" (*Spreadsheet 2*), "GSE1- bound regions only upon MC2580" (*Spreadsheet 3*) and "GSE1- bound regions that display reduced binding upon MC2580" (*Spreadsheet 4*).

**Table S6:** List of primers used for the RT-qPCR (*Spreadsheet 1*) and the ChIP-qPCR (*Spreadsheet 2*) analyses.

### References


1 Ravasio R, Ceccacci E, Nicosia L, Hosseini A, Rossi PL, Barozzi I *et al.* Targeting the scaffolding role of LSD1 (KDM1A) poises acute myeloid leukemia cells for retinoic acid-induced differentiation. *Sci Adv* 2020. doi:10.1126/sciadv.aax2746.

2 Zhang X, Smits AH, Van Tilburg GBA, Ovaa H, Huber W, Vermeulen M. Proteome-wide identification of ubiquitin interactions using UbIA-MS. *Nat Protoc* 2018. doi:10.1038/nprot.2017.147.

3 Kingston RE, Chen CA, Rose JK. Calcium Phosphate Transfection. *Curr Protoc Mol Biol* 2003. doi:10.1002/0471142727.mb0901s63.

4 Chen EY, Tan CM, Kou Y, Duan Q, Wang Z, Meirelles G V. *et al.* Enrichr: Interactive and collaborative HTML5 gene list enrichment analysis tool. *BMC Bioinformatics* 2013. doi:10.1186/1471-2105-14-128.
